# Effects of Plastic Ingestion on Blood Chemistry, Gene Expression and Body Condition in Wedge-Tailed Shearwaters (*Ardenna Pacifica*)

**DOI:** 10.1101/2022.11.26.517527

**Authors:** Nicole Mejia, Flavia Termignoni Garcia, Jennifer Learned, Jay Penniman, Scott V. Edwards

## Abstract

Plastic pollution is a global threat and affects almost every marine ecosystem. The amount of plastic in the ocean has increased substantially over the past decade, posing a mounting threat to biodiversity. Seabirds, typically top predators in marine food chains, have been negatively affected by plastic pollution. Here we focused on documenting the sublethal effects of plastic in Wedge-tailed Shearwaters (*Ardenna pacifica*, WTSH) on the island of Maui, Hawai’i. Through analyses of blood chemistry, gene expression, morphometrics and stomach contents, we documented the effects of plastic ingestion on adult WTHS from 3 established colonies. We detected a negative relationship between body weight and the presence of plastic in regurgitated stomach contents. Genes associated with metabolic, biosynthetic pathways, inflammatory responses and ribosome function were upregulated in lighter birds. Birds that had ingested plastic tended to be lighter in weight, in comparison to birds that did not have plastic and tended to weight more. Furthermore, there were 43 genes differentiating males and females that did not have plastic compared to only 11 genes differentiating males and females that had ingested plastic. There was also a marginal negative relationship between lighter birds and blood urea nitrogen levels. We also hope that the morphometric measurements, blood parameters and gene expression data we collected contributes to a database that will be used for future studies on understanding anthropogenic effects on seabird body condition.

## INTRODUCTION

Plastics are the most common form of marine debris. It is estimated that in 2010, approximately 8 million metric tons of plastic entered the ocean (NOAA, 2022). This number has continued to grow exponentially over the past decade as the worldwide production has increased nearly 200-fold since the 1950s (Ritchie & Roser, 2018). The composition of plastic products leads to detrimental environmental consequences because, unlike plant products, they do not biodegrade quickly. Estimates of plastic decomposition range up from decades to several hundreds of years, causing accumulation of plastic in the natural environment (Barnes et al., 2009).

### Plastics in the environment

Plastic has several properties that cause it to amass toxins on its surface through physical interactions such as physisorption and non-covalent bonds (Verla, 2019). Microplastics have a low-density and can be found on the surface microlayer of the ocean where they interact with heavy metals and organic chemicals. They also serve as good sorbents for heavy metals and organic chemicals due to a large surface area-to-volume ratio and hydrophobic surfaces. The substantial amount of toxins on the surface of microplastics can serve as vectors for transport of toxins to organisms (Koelmans et al., 2016).

One of the main pathways through which microplastics enter the environment is ingestion by organisms (Gallo et al., 2018). Approximately 43-100% of the world’s marine mammals, seabirds, and turtle species are at risk of ingesting plastic (Lavers et al., 2019). Ingestion endangers these organisms to perforations, entanglements, and other non-visible effects. Many marine organisms ingest microplastics through filter- or deposit-feeding, mistaking them for prey when foraging or by consuming prey that have ingested microplastics (Gallo et al., 2018).

Studies suggest that ingested plastics may cause harm to an organism because of their ability to break down in the body and leach toxins into the bloodstream. There was marked bioaccumulation of the toxins in the digestive glands and gills of mussels and lugworms that were exposed to microplastics contaminated with toxins (polyaromatic hydrocarbons, nonylphenol and phenanthrene, Browne et al., 2013, Gallo et al., 2018). Research shows that exposure to these chemicals disrupt endogenous hormones, a process that can cause reproductive complications (Gallo et al., 2018). Toxins absorbed by plastic have also been linked to neurological or behavioral changes in organisms (Gallo et al., 2018). A study that aimed at assessing the effects of polyethylene microplastics in amphibians, exposed *Physalaemus cuvieri* to 60 mg/dL of polyethylenes for just 7 days and reported visible mutagenic (da Costa Araújo et al., 2020). Other effects included accumulation of polyethylene in the gills, gastrointestinal tract, gastrointestinal tract and in the blood as well as several external morphological changes (da Costa Araújo et al., 2020).

It is acknowledged that plastic pollution has more negative effects on seabird health when compared to other marine vertebrates (Thiel, 2018). Examples of negative effects of plastics on marine vertebrates include nutritional deprivation, reduced body mass, reduced appetite and damage or obstruction to the gut (Wang et al., 2021). In a study conducted in Lord Howe Island, Lavers *et al*. (2019) found that plastic ingestion had “significant negative effect(s) on bird morphometrics and blood calcium levels and a positive relationship with the concentration of uric acid, cholesterol, and amylase” (Lavers et al., 2019). Despite these findings, few experiments have further examined sublethal effects of plastic on physiology, gene expression and overall seabird health and living species. There is a limited understanding of the effects of microplastic exposure on gene expression, particularly in vertebrates. However, experiments have shown possible effects of exposure on gene expression in Zebrafish (*Danio rerio*). Disruptions in reproduction were shown in breeding groups of Zebrafish that were exposed to environmentally significant concentrations of Bisphenol - A, a chemical used in plastic production, for 15 days (Liang et al., 2016). More studies are needed on natural populations to understand the mechanistic connections between plastic pollution and gene expression in the affected organisms.

In this study, we focused on detecting possible effects of microplastic exposure on gene expression, morphometrics, and blood analytics in three established colonies of Wedge-tailed Shearwaters (*Ardenna pacifica,* WTSH, Fig.1A). WTSH, highly pelagic seabirds that range across the tropical and subtropical areas of the Pacific and Indian Ocean, engage in feeding behaviors such as contact dipping and surface-seizing (Adams et al., 2020). It is predicted that WTSH and other seabirds with similar feeding behavior are susceptible to ingesting the floating pieces of plastic on the surface of the ocean (Boersma & Groom, 1993). Plastic ingestion by WTSH in Hawai i has been documented; however, sub-lethal effects were not evaluated (NOAA). To evaluate possible sub-lethal effects of plastic ingestion in WTSH, we collected morphometric data and blood samples to measure the health of the seabird. Blood chemistry can be used as an indicator for overall health and morphometric data are widely used in ecological studies to determine body condition (Harr, 2002, Mallory et al., 2010, Labocha & Hayes, 2012). While ornithologists debate over which morphometrics provide the best estimates for body condition, literature suggests gathering multiple proxies to fully understand body condition in seabirds (Mallory et al., 2010, Labocha & Hayes, 2012). Along with collecting several morphometric measurements such as weight and blood chemistry, we used transcriptome analysis to characterize activity of actively expressed genes under various conditions. More specifically, transcriptome analysis provided a mechanism to compare snapshots of which genes were turned on or off in seabirds that had ingested plastic and those that did not have ingested plastic. Knowledge gaps and questions remain about the intrinsic aspects of plastic, severity of impact on human health and marine organisms, effective mitigation measures, and biomagnification across the food webs (Bonanno & Orlando-Bonaca, 2018, Galloway, 2020). Therefore, the aim of this project is to provide more information to this overarching question of how plastic debris affects marine organisms.

**Fig. 1.**
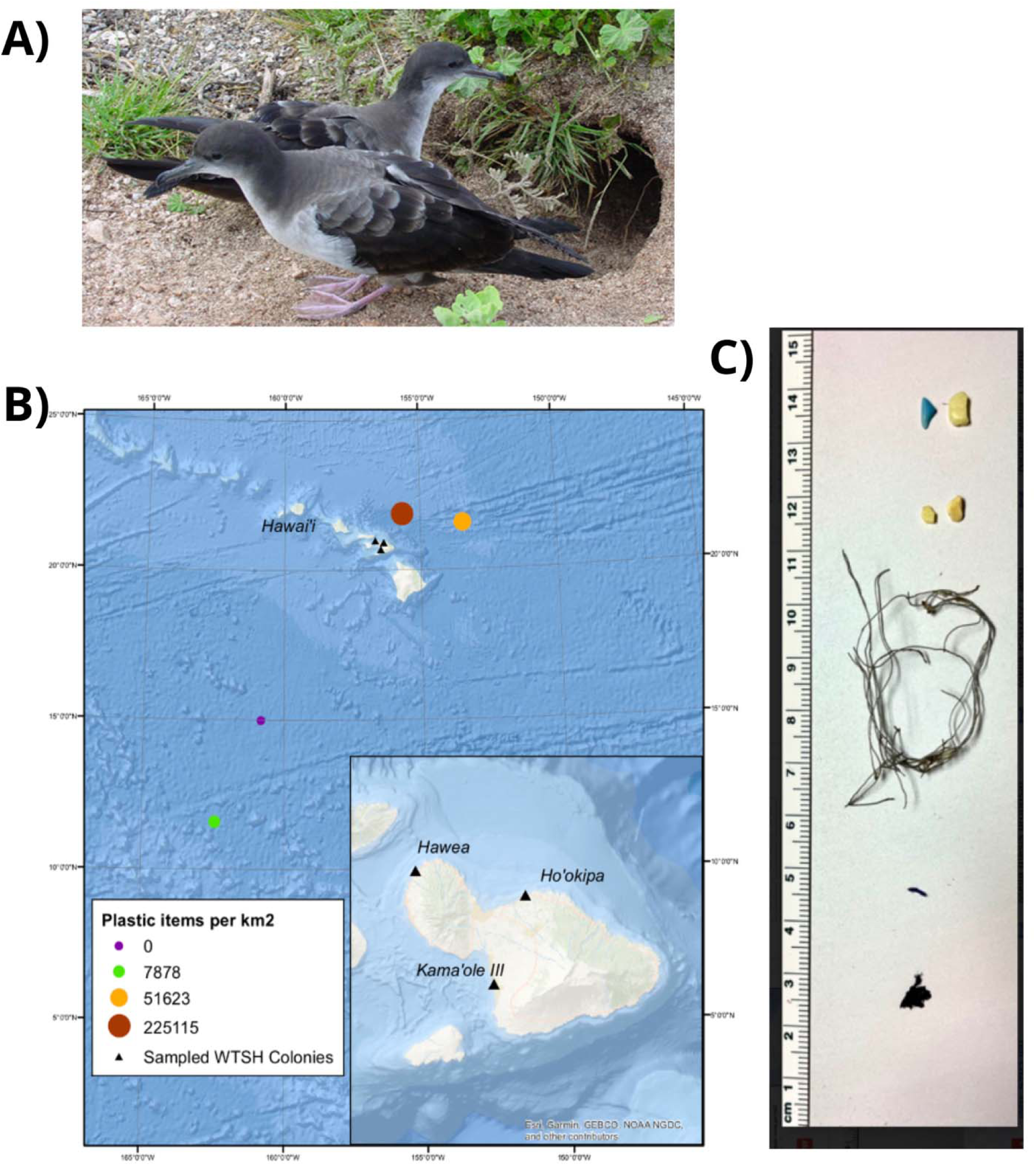
(A) Wedge-tailed Shearwater *(Ardenna pacifica)* in Hawai’i. From U.S. Fish and Wildlife Service by Ian Jones. (B) Sampling Sites. Three sampling sites on the island of Maui in the Hawaiian Islands represented by triangles. Colored circle represents concentrations of plastic (items km-2) found around the Hawaiian Islands based on data from Cózar et al. 2014). (C) Plastic samples. Plastic collected from gut samples from WTSH.

## Materials and Methods

### Study site

The collection of blood samples occurred at three different sites on the island of Maui, Hawai’i. These sites were chosen based on known established colonies of WTSH at these locations (Fig. 1B). Although all populations are located in protected areas, the three sites were on the seashore near frequently visited beaches. Kamaole Park III is located on the southern part of the island. Ho’okipa Beach located on the northern part of the island. Finally, Hawea Point is located on the western part of the island; this location also had the largest WTSH colony.

These populations of WTSH along with other seabird populations are vulnerable to microplastic exposure given that there are significant centers of plastic accumulation around the Hawaiian archipelago. Andrés Cózar et al. (2014) synthesized data to create a global map that approximates the magnitude of plastic pollution in the open ocean. Using the coordinates and data from Cózar et al. (2014), we mapped the magnitude of plastic pollution near the island of Maui (Fig. 1B). The concentration near Maui is sizeable compared to other sampled locations, with a non-adjusted concentration of 225,115 items per km^2^. Despite our collection of samples from three different locations, we do not expect any significant genetic differentiation between the populations. Whereas marked genetic differentiation has been found in WTSH in populations breeding in different islands and archipelagos along the Pacific coast of North America, there is no evidence for deep evolutionary divergences (Herman et al., 2022). The three WTSH colonies were treated as one population when we analyzed the data.

### Gut Sample Testing

We used the procedure outlined by Duffy et al. (1986) for gut sample testing to collect possible ingested plastic. The procedure outlined by Duffy et al., involves a stomach pump system where the seabird is filled with seawater through a gavage and then tipped over a bowl to promote and collect regurgitation. We completed sampling of the gut contents for 28 out of the 29 birds.

### Blood Sampling

We collected blood samples first to avoid a signal of stress response in the analyses of gene expression and blood chemistry from being handled. Blood sample collection was possible in 28 out of the 29 birds for gene expression analysis, and in 25 out of the 29 birds for chemical analysis. This lack was due to not enough blood collection during sampling.

We collected approximately 200µ l of blood using a syringe from the medial metatarsal vein. We used styptic powder to stop bleeding when necessary. We added 100µ l of blood to a vial containing RNAlater buffer. We stored 20µ l of blood in heparin tubes for iStat cartridge analysis. The iStat Chem8+ provided us with the following blood analytics; sodium (Na mmol/L), potassium (K mmol/L), chloride (Cl mmol/L), ionized calcium (iCa mmol/L), total carbon dioxide (TCO2), glucose (Glu mg/dL), urea nitrogen/urea (BUN mg/dL), creatinine (Crea mg/dL), hematocrit (Hct %PCU), hemoglobin (Hb g/dL), anion gap (AnGap mmol/L). We used Qiagen QiAMP DNA Blood kit for DNA purification to proceed to the PCR reaction. We used the universal method outlined by Fridolfsson and Ellegren (1999) for sexing in birds with PCR reaction. The 2- primer system is as follows:

2550F: 5’-GTTACTGATTCGTCTACGAGA-3’
2718R: 5’-ATTGAAATGATCCAGTGCTTG-3’

Using this primer system, we employed standard PCR on the templates of DNA extracted from unknown-sex *A. pacifica* species. The PCR mixture (15 µ l) contained 1.5 µ l of 10X buffet, 0.5 µ l of dNTP (10 pmol), 0.5 µ l of forward primer (10 pmol), 0.5 µ l of reverse primer (10 pmol), 0.1 µ l of NEB Ta1 (5U/uL), and 9.4 µ l of H_2_O. 2.5 µ l of the DNA extraction was used. The PCR program was as follows: 94°C for 5 min, 94°C for 30 sec, 60°C for 30 sec *touchdown, –1.0°C/cycle x 10 cycles, 72°C for 30sec, 94°C for 30 sec, 50°C for 30 sec x 30 cycles, 72°C for 30 sec, 72°C for 5 min, 4°C hold. We used molecular graded H_2_O as a negative control. A negative control was essential for possible misinterpretation due to contamination or other factors. We separated the PCR product through electrophoresis on a 2% agarose gel at 90 V for about 1 hour. We stained the gel with *GelRed*™ - a fluorescent nucleic acid gel stain that replaces the highly toxic ethidium bromide (EtBr) - and we used a gel imaging camera.

### Morphometric Data

We completed collection of morphometric measurements for 28 out of 29 of the birds, using a Pesola scale. We measured tarsus length, bill length, nares depth and width using calipers and we measured wing chord length using a ruler. We banded each bird and recorded the number if the bird was a recapture.

### RNA Isolation and Sequencing

Before initiating RNA isolation, we removed RNAlater through centrifuging aliquots at 20,800 x g (RCF). We removed supernatants from the remaining pellets containing cell material. We used a Qiagen RNeasy Plus Universal Mini Kit for blood isolation. At the Harvard Bauer Sequencing Core Facility, we used KAPA mRNA Hyperprep kit and a NOVASeq SP platform to sequence paired end reads of 150 bp length, yielding between 20 and 30 million reads per sample.

### Data analysis

We calculated several values to assess the quality of the RNA. We analyzed RNA integrity score (RIN) values, and calculated Phred Scores to assess the quality of our sequencing. RIN values assign a numerical value to the quality of the RNA that we worked with to evaluate the integrity of 18S and 28S rRNSs (Puchta et al., 2020). A RIN value of 8 and above indicated higher quality and integrity of RNA and values below 5 indicated some levels of RNA degradations. Phred scores are similar in that they assess the quality of sequences. Similar to RIN value, a high Phred score (90% and above) indicated better quality sequences (Scholz, 2021).

We aligned sequences to the publicly available reference genome of Cory’s Shearwater (*Calonectris borealis,* accession # PRJNA545868, Feng et al., 2019) with the RNA sequence mapper STAR (Spliced Transcripts Alignment to a Reference) (Dobin et al. 2013). Followed by a transcript quantification with RSEM (Li & Dewey, 2011) and a differential gene expression analysis with DESeq2 (Love et al., 2014) in R programming. We created heatmaps to determine if there were observable patterns between gene expression and presence of plastic. Combined with clustering methods, heatmaps can help determine there are similar changes in gene expression based on their activity. In all of the tests, we used both a conservative p-value of 0.05 and a less conservative p-value of 0.1 to determine the significance of results. The summary of all tests we ran can be seen in supplementary table 1.

We conducted gene ontology analysis using R 3.5.1 (R Core 226 Team 2018) and the package ggprofiler2 with *Gallus gallus*, *Taeniopygia guttata* and *Mus musculus* as the model systems in the search database. We used Ggplot2 and plotly for plotting as outlined in Kolberg et al. (2021). We separated terms into Gene Ontology, KEGG pathways and Reactome databases.

We ran multiple tests to analyze possible relationships between the variables of sex, presence of plastic and blood chemistry levels. We ran all analyses using R 3.5.1 (R Core Team 2018). We used a t-test to compare the means between blood analytes of birds that had ingested plastic and those that had no. All blood parameters were analyzed independently. Parameters were plotted against the presence of plastic. Differences were considered statistically significant when p < 0.07. We ran principal component analysis (PCA) to determine if the categories of plastic and sex where clustering together according to the morphometric, blood chemical and genetic data. We first ran PCA in relation to sex to control for possible differences in sex. We then ran PCA in relation to the presence of plastic to determine if individuals would cluster due to physiological differences caused by the presence of plastic. Packages used for these analyses included devtools, ggplot2 and ggbiplot. Finally, we used a general linear model (glm) to study the association between the variables, such as the morphometric measurements and blood parameters, and the conditions, which was either presence or absence of plastic. Differences were considered statistically significant when p < 0.05. and we considered effect size as well.

## Results

### Gut samples

Plastic or other unidentified hard pieces were found in 12 of the 28 birds sampled for plastic (Fig. 1C). These included fishing line and pieces of microplastics. A summary of which birds contained plastic, their sex and what blood samples were available for each bird can be found in Fig. 2. Four out of the ten males contained plastic or other unidentified hard pieces. Seven out of 17 females contained plastic or other unidentified hard pieces. Bird N004 contained plastic or other unidentified hard pieces, but the sex of this bird was unknown due to insufficient blood sample to carry out analyses (Fig. 2).

**Fig. 2.**
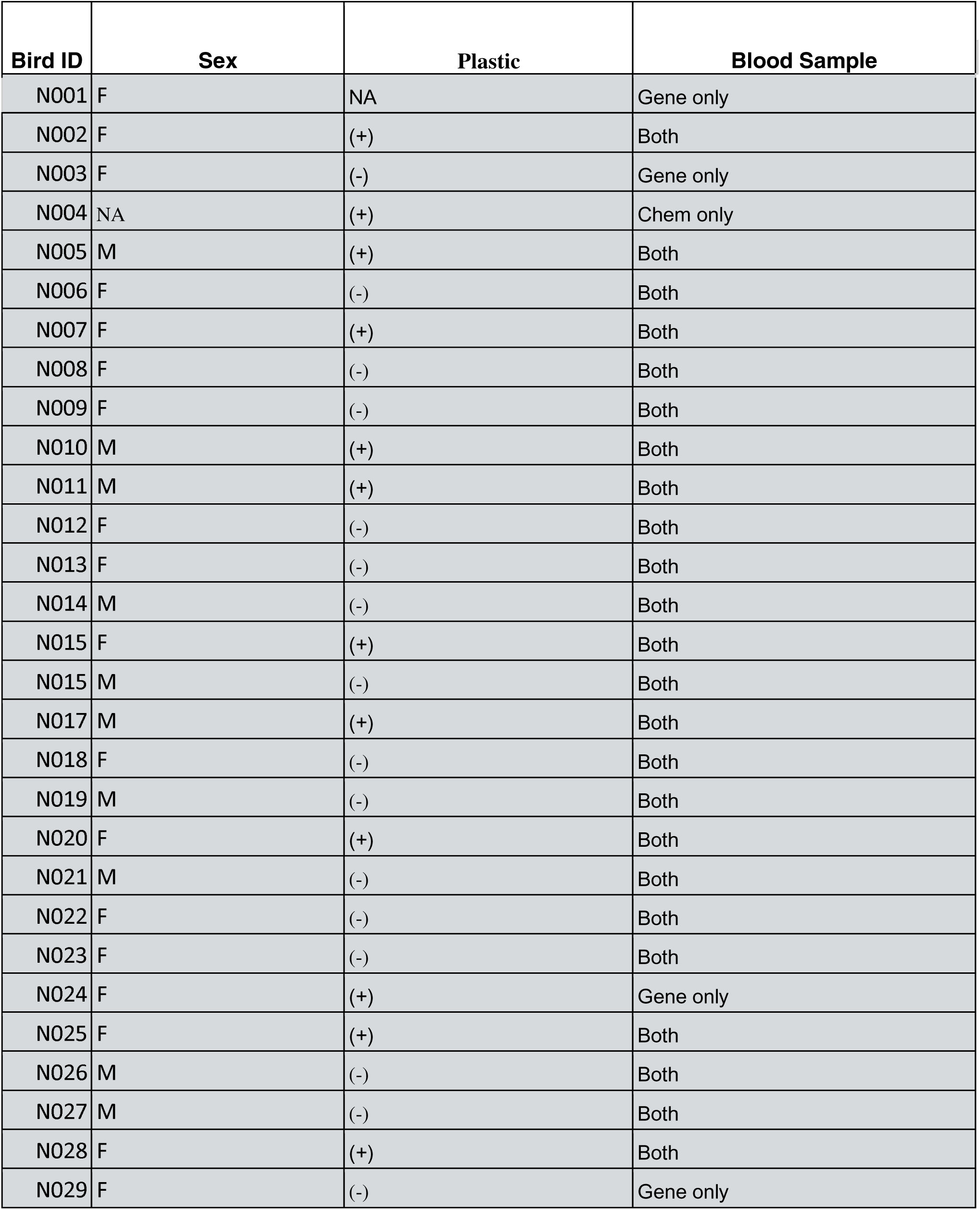
Bird ID and data available for each specimen. An ID was appointed for each specimen. The sex is indicated for each specimen. Plastic refers to if that particular specimen had ingested plastic; (+) indicates presence of plastic and (-) indicated that that specimen did not have ingested plastic. Blood sample refers to what information was obtained from the collected blood. Specimens where only genetic information is available is denoted by “Gene only”. Specimens where only a chemical blood panel is available is denoted by “Chem only”. Specimens where both genetic information and chemical blood panels are available is denoted by “both”.

### Sex Determination through PCR

Our sample consisted of 10 males and 17 females. Bird N001 was a female, but gut samples were not collected for this bird (Fig. 2).

### Blood Analytics, Morphometric and Gut Sampling

We found minimal relationships between the presence of plastic and the measurements listed above. Significant values from t-test analyses included levels TCO2 (Fig. 3e.) in the presence of plastic. There were no other significant values for other blood analytes in our t-tests and t-values ranged from .05789 to .87 (Fig. 3).

**Fig. 3.**
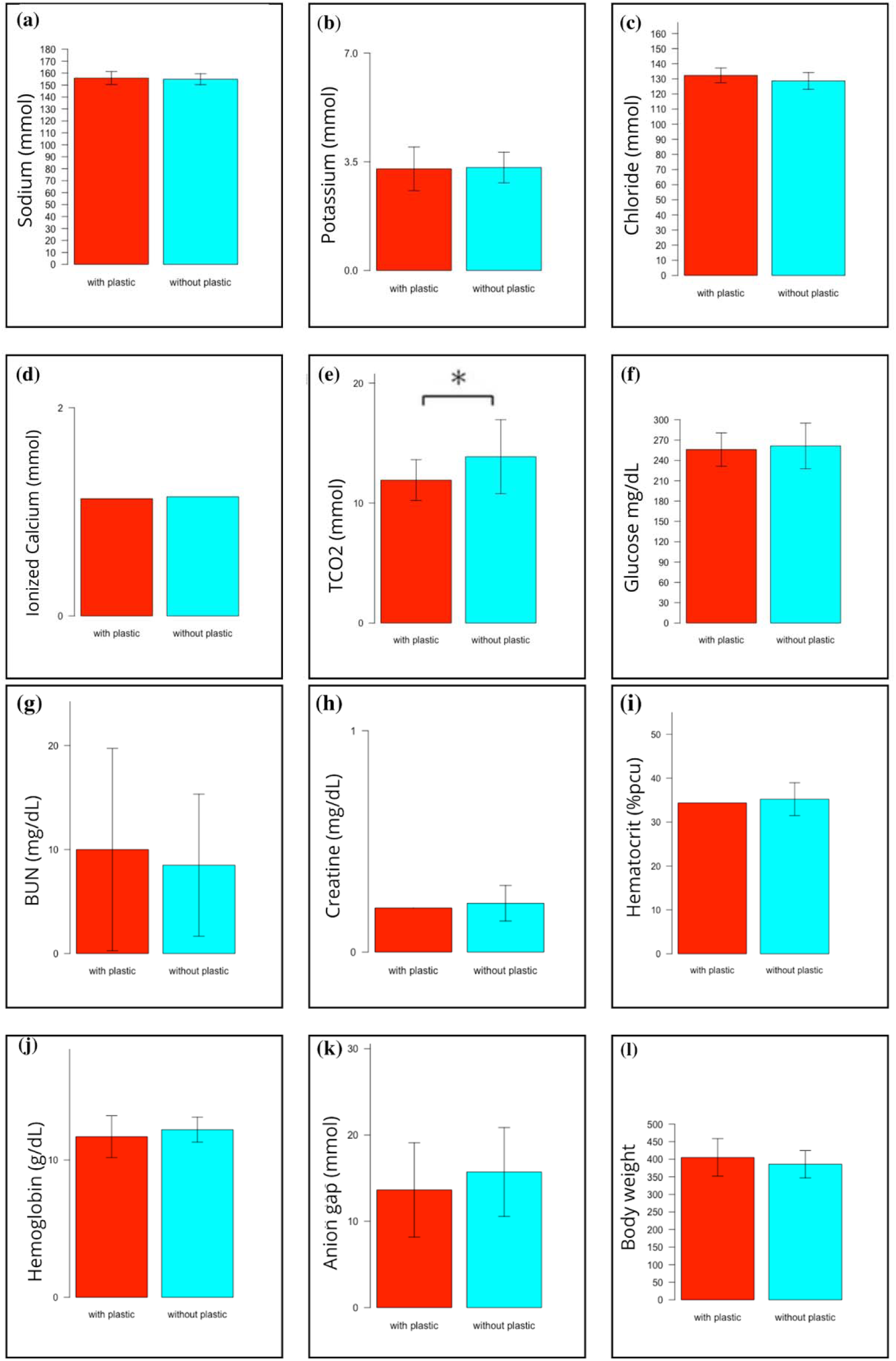
Individual t-test analysis of each blood analyte and weight. (a-k) Blood analyte plotted against presence and absence of plastic with error bars. Any test that showed a slight significance value is noted on the right-hand corner of the individual chart. (l) Weight plotted against presence or absence of plastic.

Our PCA results showed minimal relationships for both blood chemistry and morphometric measurements when the independent variable is set as either sex or presence of plastic (Fig. 4). The absence of clustering when using blood chemistry and morphometrics provides a control for sexual dimorphism in our samples. Wedge-tailed Shearwaters are not sexually dimorphic, which is consistent with our results.

**Fig. 4.**
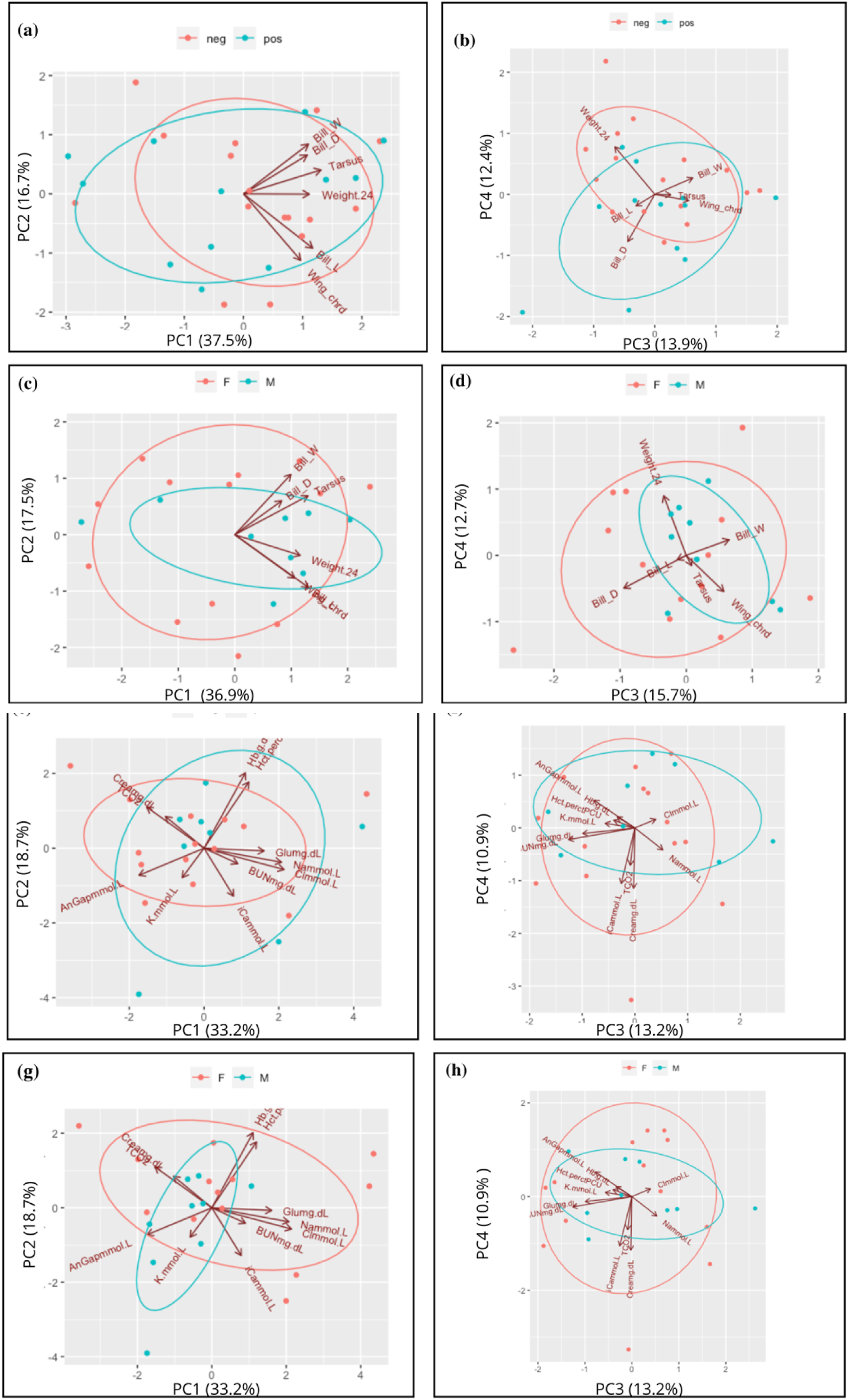
PCA analyses of morphometrics and blood analytes. (a) Morphometric measurements with variables of presence of plastic using PCA 1 and PCA 2. (b) Morphometric measurements with variables of presence of plastic using PCA 3 and PCA 4. (c) Morphometrics measurements with variables of sex using PCA 1 and PCA 2. (d) Morphometric measurements with variables of sex using PCA 3 and PCA 4. (e) Blood analytes with variables of presence of plastic using PCA 1 and PCA 2. (f) Blood analytes with variables of presence of plastic using PCA 3 and PCA 4. (g) Blood analytes with variables of sex using PCA 1 and PCA 2. (h) Blood analytes with variables of sex using PCA 3 and PCA 4.

Weight deviated from the other variables in the presence or absence of plastic under PCA 3 and 4 (Fig. 4b). There was a negative relationship between weight and presence of plastic when using a general linear model; birds that had ingested plastic tended to weigh less whereas birds that did not have ingested plastic tended to weigh more (Fig. 5a). When weight was the independent variable, urea nitrogen/urea, hematocrit and potassium demonstrated significant p- in Supplementary figure S2.

**Fig. 5.**
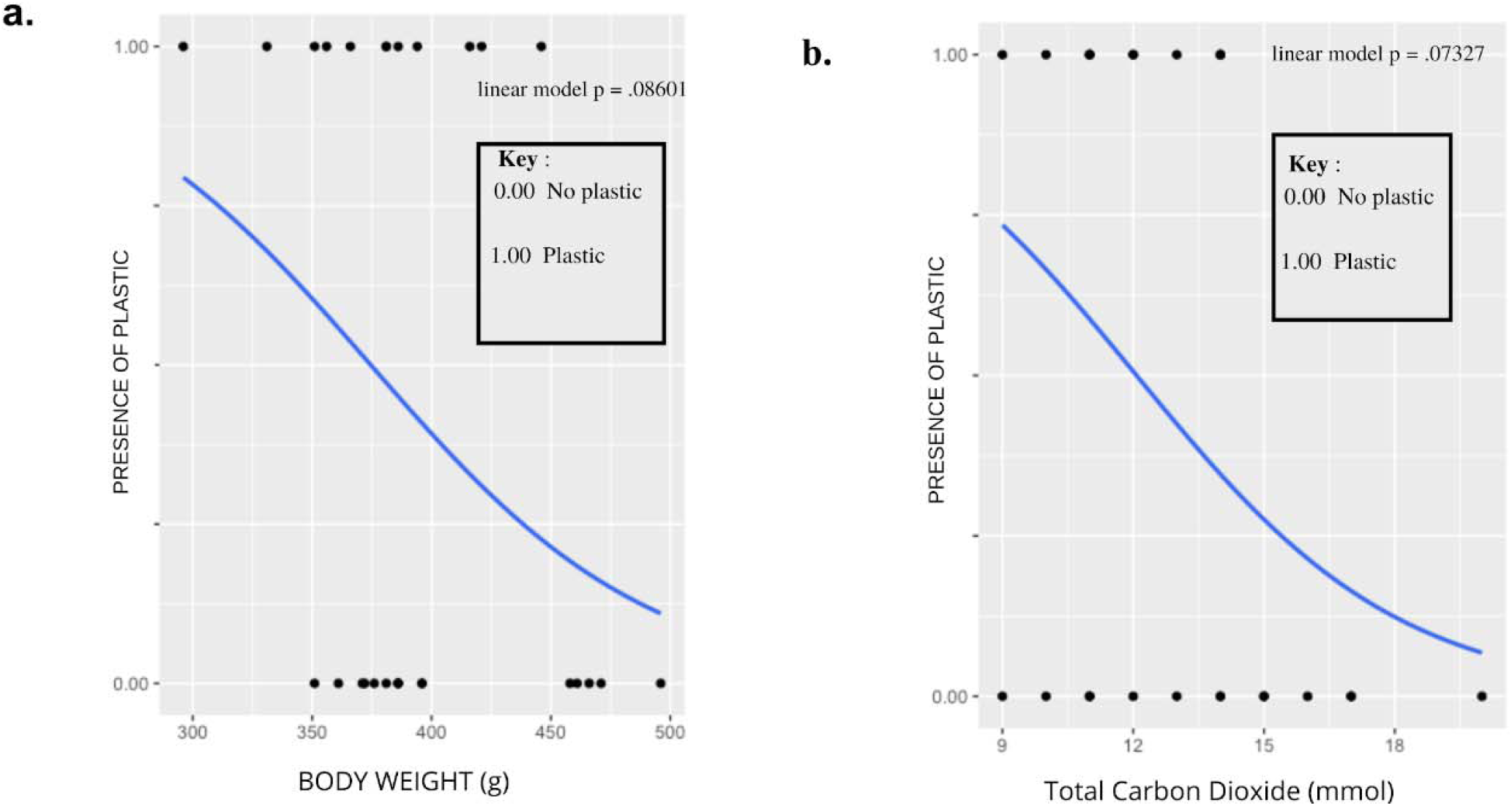
Significant relationships from general linear model. (a) Relationship between presence or absence of plastic and weight of bird. (b) Relationship between presence or absence of plastic and total carbon dioxide.

**Fig. 6.**
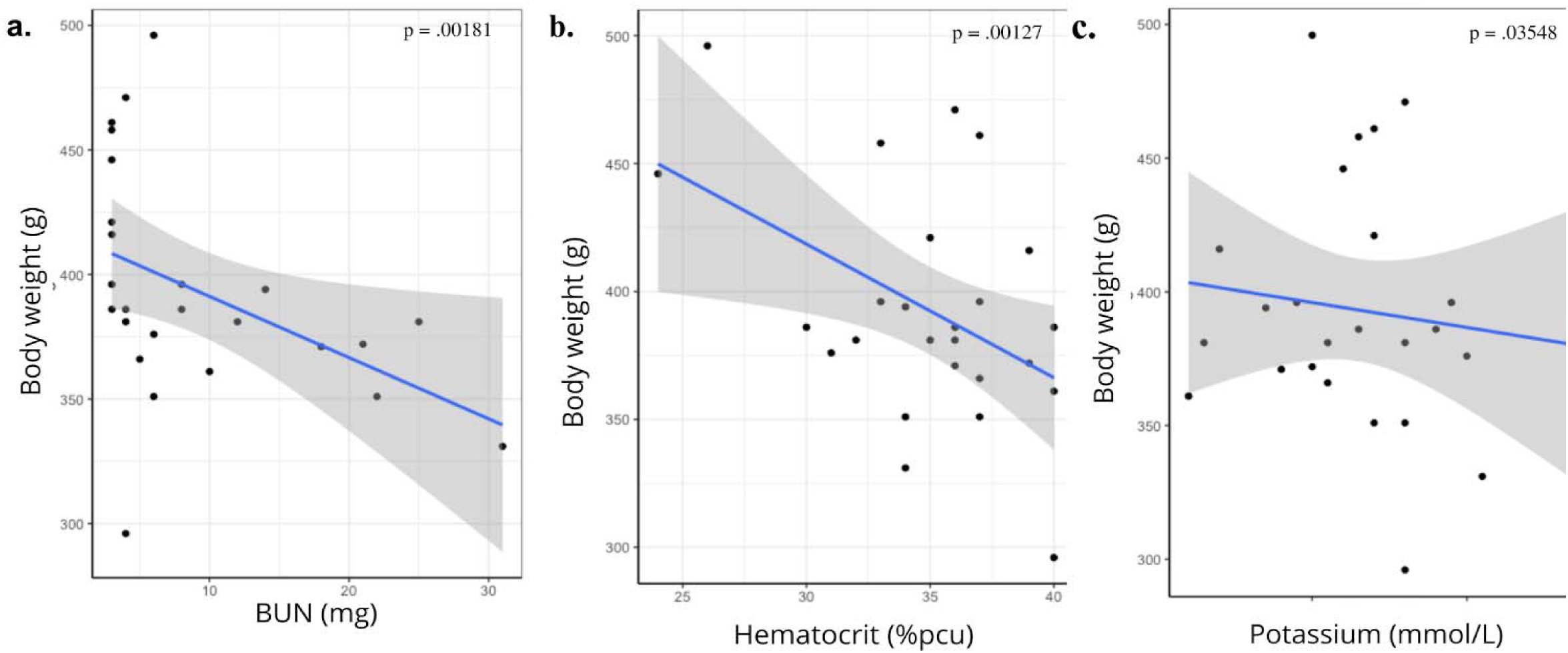
Significant results from general linear model of blood analytes with weight as the variable. (a) Hematocrit as percentage with weight as variable. (b) Urea nitrogen/urea with weight as the variable. (c) Potassium with weight as the variable.

### R NA Isolation and Sequencing

The RIN value used to assess RNA integrity during isolation was around 8.4 - 6 for most samples (Supplementary figure S9). Sample N005 had a lower RIN value of 4.8. Phred Scores were used to assess sequencing quality (Supplementary figure S4). On average, all samples reached > 30 Phred Score, which indicated good quality for downstream analyses. Supplementary figure S4 depicts reads per sample, which fell between 48,000,000 and 80,000,000 reads per sample.Uniquely mapped reads were all above 40%, with the highest being slightly above 60% (Supplementary figure S5). Low values could be due to the rate of degradation of blood RNA (Dobin & Gingeras, 2015) or because we are not using a species-specific reference genome for the transcriptome alignment. Multiple mapped reads, which read mapping to multiple locations in the genome, occurred at a rate of 1.3% to 2.7% (Supplementary figure S6, Pantano, 2018). We were able to distinguish four genes that separate the samples into two groups, discernible by upregulation and downregulation activity. Shades of blue represent downregulation activity while shades of red indicate upregulation of the gene (Fig. 7A).

**Fig. 7.**
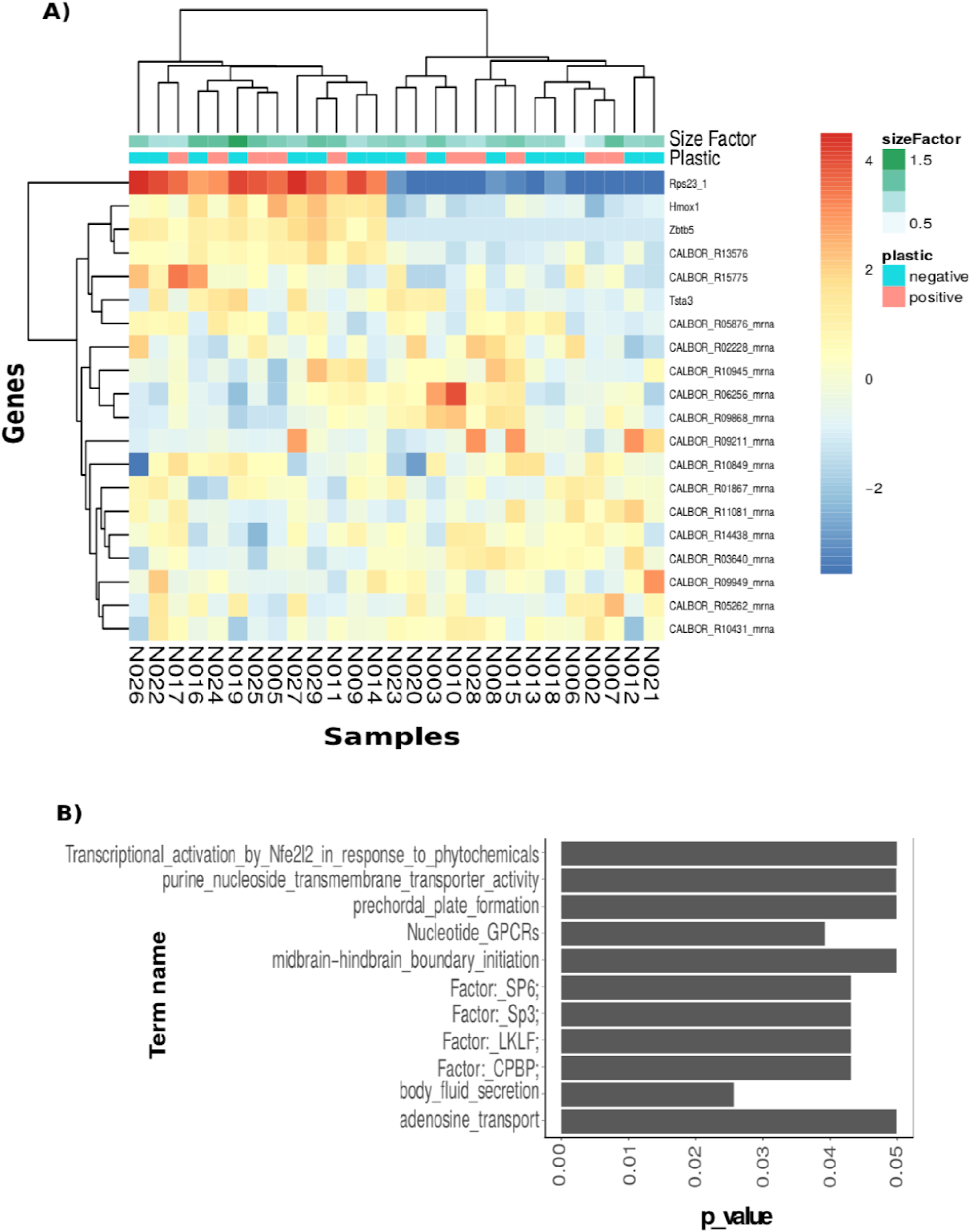
DE genes and enrichment analysis for the entire data set. (A) Heatmap showing that plastic and size factor of libraries do not explain the DE gene profiles. The heatmap shows the top 20 genes with the highest statistical power. (B) Enrichment analysis and significant terms for the 6 DE genes shown on the heatmap above.

There were no discernible differences in gene expression under the presence and absence of plastic. These results suggested that the clustering by presence of plastic does not explain the differential gene expression profiles (see Fig. 7A). We did not obtain significant results with either the conservative p-value or less conservative p-value.

We divided the analysis into sex; females in one analysis and males in the other with and without plastic as the variable (Supplementary figure S7). In this analysis we identified one gene that differentiated females with and without plastic when using the less conservative p-value. Fourteen genes differentiated males with and without plastic under the more conservative p-value. The presence of plastic in males was associated with the upregulation of the top four genes. Eleven genes differentiated males and females that contained plastic (Supplementary figure S8A). Forty-three genes differentiated males and females that did not have plastic (Supplementary figure S8B).

We analyzed DE genes that examined weight with and without plastic. We divided weight into 3 factors: low, medium and high. Birds that did have ingested plastic, tended to be heavier and showed a downregulation in the expression of the top 18 differentiating genes (Figure 8A). Birds that had ingested plastic, tended to be lighter and showed an upregulation in the expression of the same genes. These distinctions were able to be made while using a conservative p-value for the top four genes and marginal p-value for the remaining genes. The genes responsible for differentiation when the samples were not separated into variables were associated with transcriptional activation, body fluid secretion and protein transportation activity (Fig. 7B). The top two genes responsible for weight differentiation, Ankrd11_1 and Hsph 1 (Supplementary figure S10A), were upregulated in heavier birds. Genes that were upregulated in heavier birds were associated with trimethylation and cell cycle function (Supplementary figure S10B). The top twelve genes that were upregulated in lighter birds (Fig. 8A) have been associated with several metabolic and biosynthetic processes, ribosome function and pathway response associated with COVID-19 (Supplementary figure S10).

**Fig. 8.**
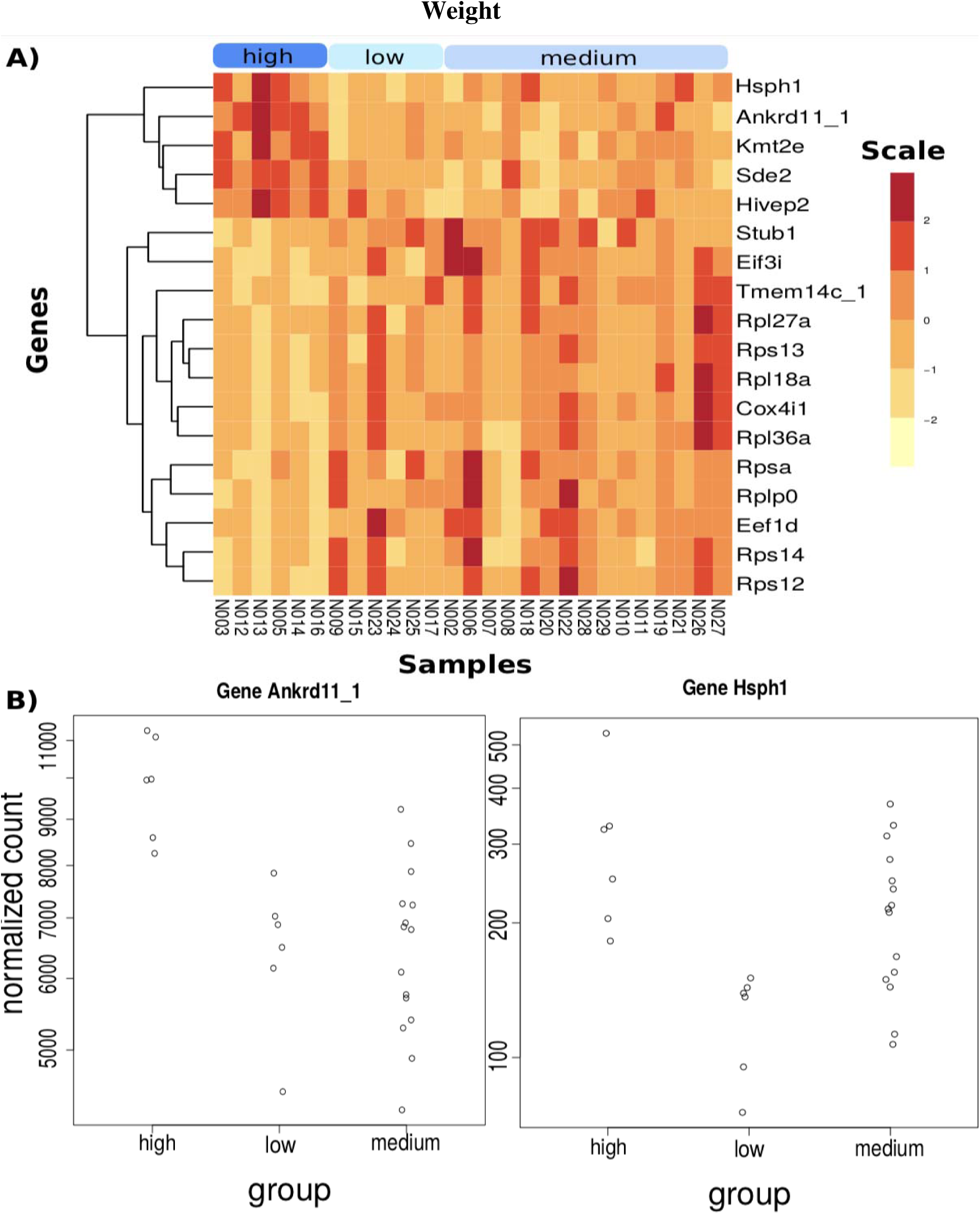
DE genes with three categories of weight; low, medium and high. (A) Heatmap showing the 18 significantly DE genes. (B) Normalize count in the two top genes showing differences in counts between the three weight categories.

## DISCUSSION

Our overarching question was whether ingested plastics from the environment have a sublethal effect in seabirds, gut samples, morphometric measurements and blood samples from Wedge-tailed Shearwaters in Maui, Hawai’i as a test case. Using PCA analysis, general linear models and analysis of DE genes, we quantified the effect of microplastic load on the overall health of the birds. We found a marginal, negative correlation between body weight and plastic. When using body weight as an indirect measure for the effect of plastic on the bird, we found associations between upregulation of metabolic and biologic pathways and lighter birds. Birds with lower weight tended to contain plastic.

The main effects to gene expression were attributed to i) upregulation of biosynthetic and metabolic pathways in lighter birds and ii) downregulation of biosynthetic pathways in heavier birds. Analysis of DE genes showed an upregulation of genes involved in biosynthetic processes in lighter birds. Biosynthetic and metabolic processes are responsible for body mass accumulation. Disruptions to energy and lipid metabolism pathways have been documented in Zebrafish (*D.rerio*) (Limonta et al., 2019, Lu et al., 2016), and African Catfish that were exposed to microplastics (*Clarias gariepinus*) (Karami et al., 2016). Toxins found in microplastics, such as BPA and BP, are associated with disruption of metabolic and biosynthetic pathways (Sun et al., 2022). Sprague-Dawley Rats exposed to BPA experienced weight loss (Sun et al., 2022). Adult Zebrafish, also exposed to BPA, experienced a decrease in oxidative stress and body weight change (Jordan et al., 2012, Lu et al., 2014, Huang et al., 2019, Sun et al., 2022). Previous physiological suggestions in seabirds include reduction in the functional volume of the gizzard; this reduces digestive capability causing weight loss (Furness & Monaghan, 1987).

Lighter birds also displayed upregulation of genes involved in organonitrogen compound metabolic processes. This process is associated with the formation of compounds that are directly linked to a nitrogen atom. Our general linear model revealed a marginal relationship between lighter birds and higher BUN (blood urea nitrogen) levels. Higher BUN levels have been used to infer dehydration in birds (Hochleither). Higher urea levels were associated with European pond turtles (*Emys orbicularis*) exposed to varying doses of microplastics (Banaee et al., 2020). These results could suggest a possible relationship between the presence of plastic, and disruption of genes involved in metabolic and biosynthetic processes.

There were only 11 genes that differentiated females and males among individuals with plastic (Supplementary figure S8A). Meanwhile, there were 43 genes that differentiated females and males among individuals without plastic (Supplementary figure S8B). Several studies have found that when bisphenol-a (BPA), a chemical used in plastic, is leached from plastic, it can activate estrogen receptors in mammals (Bittner et al., 2009, Gao et al., 2015). These findings suggest that males would present a detectable level of estrogen, causing the genetic differentiation to be less defined between males and females. More research is needed on the effects of plastic toxins on sex hormones to support this hypothesis.

One of our significant findings included genes associated with a pathway in response to a COVID-19 infection (Supplementary figure S10B). This pathway is associated with an inflammatory response, organ failure, and hypercoagulability (Harrison et al., 2020). We cannot be sure that this has to do with a Sars-CoV-2 infection in birds, but this pathway response in birds may be associated with a similar infection leading to an inflammatory response. Male mice exposed to PCBs, DDE (dichlorodiphenyldichloroethylene) and HCB (hexachlorobenzene), suffered liver injury and systemic inflammation (Deng et al., 2019). PCBs, DDE and HCB are chemicals found in plastic. Presence of microplastics in zebrafish have been linked to enhanced immune responses (Limonta et al., 2019), and other disruptions to the immune system (Powell et al., 2010).

### Challenges in study design

One significant challenge in our study was the possibility of false negative results. All bird populations we sampled are on the island near a big source of plastic. It could be the case that those birds without plastic in their gut at the moment of sampling, were already exposed at some level to plastic and we were not able to detect it. The flushing technique for gut sampling we used does not allow for full collection of gut samples. Other methods of obtaining sampling might be more effective but could also increase the risk of death and injury to the birds. A study that does not rely on destructive sampling also allows for repeated sampling and continued monitoring of bird health. Destructive sampling only allows for a single observation (Provencher et al., 2019).

### Using weight as an indirect metric

With a marginal p-value, lighter birds tended to be associated with the presence of plastic while heavier birds tended to be associated with the absence of plastic (Fig. 5a). More importantly, the relationships that we detected have been noted in previous studies examining plastic load and its effect on different organisms (Ryan 1986, Sievert & Sileo 1993, Pierce et al. 2004). A study on *Physalaemus cuvieri* tadpoles that were exposed to polyethylene microplastics at significant concentrations for 7 days reported accumulation of microplastics in the internal organs of which led to morphological and mutagenic changes; these changes can have effects on health and development (da Costa Araújo et al., 2020). In the same study, abnormalities were observed in nuclear erythrocytes and on external morphological traits such as mouth-cloaca distance.

Despite the challenges presented by experimental design, we were able to detect marginal indications of possible effects of microplastic toxins in relation to genetic expression, weight and blood analytes. Perhaps the challenges of conducting an ecological study in the field cause the associations to not be as strong as they could have been in a laboratory setting.

## CONCLUSIONS

Our results indicate that there are signs of sublethal effects of ingested plastic in Wedge-tailed Shearwaters. We were able to detect a negative relationship between the presence of plastic and weight. When using weight as an indirect measurement for the effect of ingested microplastics, there is some evidence that plastic affects metabolic and biosynthetic processes in Wedge-tailed Shearwaters. Whereas there was no direct relationship between load of plastic collected and DE genes, there was upregulation of genes involved in biosynthetic processes and ribosome function in lighter birds. Birds that had ingested plastic tended to be lighter. There were more genes that differentiated females and males that had not ingested plastic than females and males that had ingested plastic. Furthermore, there is the possibility of false negatives (in absence of plastic loads during gut sampling). The experiment contributes to an understanding of the relationship between plastic and sublethal effects in seabirds. Given the finding of a COVID-19 response pathway, we can further ask more questions on how anthropogenic diseases are translating into wildlife. In seabirds specifically, this raises the question of whether ingested plastic is having an impact on seabirds’ immune system.

## Acknowledgements

The research leading to the results was funded from Harvard College Office of Undergraduate Research and Fellowships, and Harvard’s Museum of Comparative Zoology.

## Data Availability Statement

The sequence data will be available upon publication. Data sets associated with this paper will be available upon acceptance.

## Supporting Information

List of Supplementary figures and Tables. Tables. Supporting Information, Tables are both in the Supporting Information, File (pdf) and in the Supporting Information.

### Supporting Text

#### Heavy metals and organic pollutants

Heavy metals, metals with density greater than 5 g/cm3, find their way into the environment both through natural means and as cause of human activity (Briffa et al., 2020). Weathering of earth’s crust, urban runoff, industrial waste, pesticides, sewage runoff and many other anthropogenic sources introduce heavy metals into the environment. Heavy metals are found in significant concentrations near areas of anthropogenic activity such as harbors and marinas (Bighiu, 2017). Conversely, these are also areas where there are significant amounts of microplastics (Claessens et al.2011). Heavy metals then attach to the surface of microplastics due to the strong physical interactions. In excess quantities, heavy metals are toxic to organisms (Furness & Monaghan, 1987).

Organic chemicals, pollutants containing carbon bonded with other compounds, include persistent organic pollutants (or POPs) (Liu et al., 2021). POP’s are resistant to degradation and can bioaccumulate to toxin levels. Bioaccumulation refers to the accumulation of a contaminant in an organism. Similar to heavy metals, POPs can be traced back to both natural and anthropogenic activities. These activities include volcanic eruptions or synthesis of chemicals. POPs are known to be easily transported from the source and easily absorbed in a new environment (Ashraf, 2017). Due to the low solubility in water, they are also easily absorbed by microplastics (Verla et al., 2019). Some well-known examples of POPs include the insecticide DDT, PCBs (Polychlorinated biphenyls) and BPAs (Bisphenol A) (Verla et al., 2019).

#### Wedge-tailed Shearwater (A. pacifica) Biology

Wedge-tailed Shearwater are pelagic seabirds that are monogamous and are known to be natal philopatric. Shearwater pairs often form a long-term pair bond which lasts several years. They have extensive feeding ranges, with a mean maximum range of 615 km (Adams et al., 2020). Although not endangered, global numbers are in decline. When it is not breeding season, Wedge-tailed Shearwaters take long migrations and use specific migratory routes that take advantage of the oceanic wind patterns (Schaffer et al., 2006) It has been documented that the birds sometimes make long dispersive movements (Weimerskirch et al. 2020).

#### Science of plastic accumulation in the ocean

Due to ocean circulation patterns, there are certain regions in the open ocean where there is a greater concern for ocean pollution, such as the Great Pacific Garbage Patch in the Northern Pacific that stretches from the west coast of North America (Cózar et al., 2014). These garbage patches form because of gyres, large systems of circulating water in the ocean. Five gyres in particular play an important role in circulating water around the globe: North Atlantic, South Atlantic, North Pacific, South Pacific and Indian (NOAA). Plastic congregates around these slow-moving whirlpools, forming massive areas of circulating plastic (NOAA).

#### Gut sampling from the proventriculus

The flushing technique empties out gut contents from the proventriculus of a bird, but we cannot be assured that it empties out gut contents from the ventriculus in Procellariids (Duffy & Jackson, 1986). Procellariids’ stomachs can be divided into two sections: the proventriculus and the gizzard. A lack of plastic content in the proventriculus often means that it is either regurgitated or emptied quicker than the gizzard (Nania & Shugart, 2021). This creates the possibility that the proventriculus of birds we sampled had been already cleared of plastic and we did not fully capture the plastic load.

#### Challenges of gene expression with blood

Using whole blood to create a genetic profile is a relatively new approach, especially in non-model organisms and livestock (Désert et al., 2016). Most gene profiles use tissue samples for a particular study because the composition and content of RNA, responsible for genetic activity that we are able to investigate, is specific to the tissue activity (Jax et al., 2018). Target tissues also provide information on specific adverse effects specially in response to toxic exposure (Lobenhofer et al., 2008).

The use of whole blood for genetic profiling is a rising and useful tool (Désert et al., 2016). It could be used as a new approach in conservation to assess the health status of natural populations of species in threatened status. Studies with whole blood transcriptome have quantified immune response in birds and identifying sex chromosome evolution in two rare species of kiwi birds (Désert et al. 2016, Ekblom et al. 2014, Ramstad et al. 2016, Sandford et al. 2012). It is worthy to note the importance of continuing to advocate for these procedures which may provide a less invasive way of conducting data collection.

**Fig. S1.**
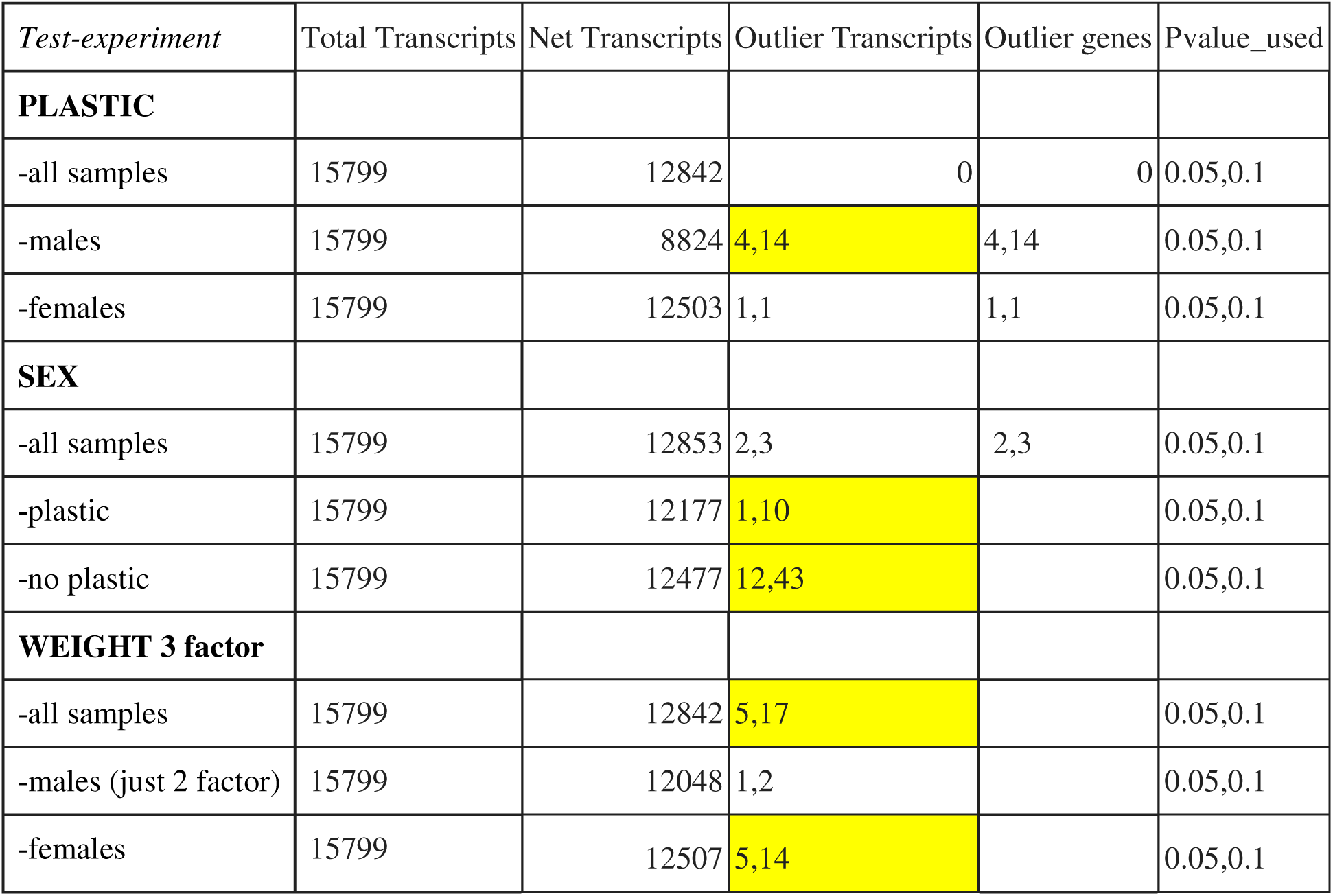
Table summarizing differential genetic expression analyses. The test/experiment column describes the three main analyses that we conducted and the variables that we used for each test. Transcripts represent the count of transcripts that were aligned with the reference genome. Net transcripts represent the count of transcripts that were represented or quantifiable in all of the samples. Outlier transcripts and outlier genes represent the count of transcripts that differentiated within the respective test. P-value represents the two p-values we used to determine significance of the results. We used both a marginal p-value (0.1) and a standard value (0.05).

**Fig. S2.**
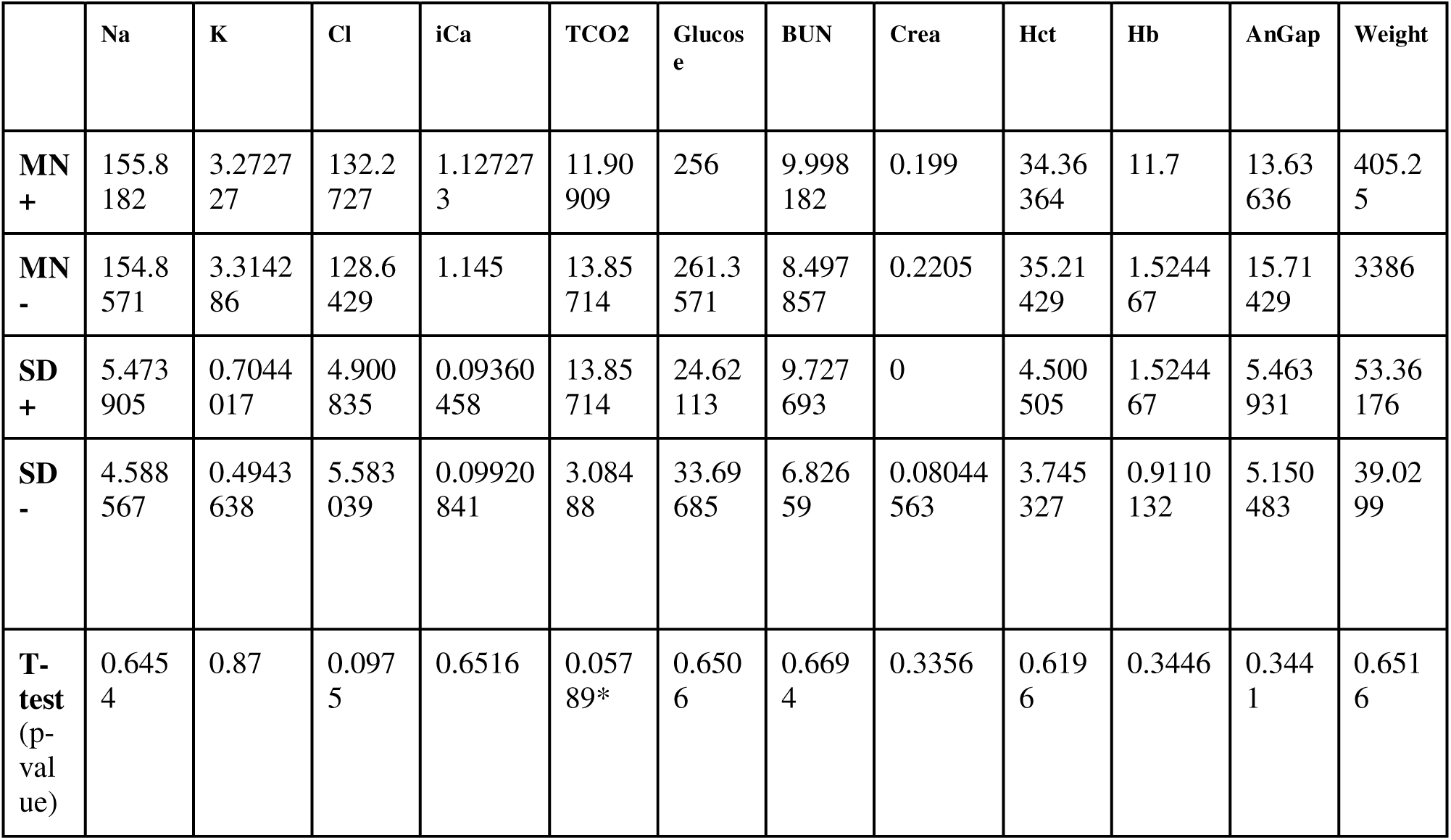
Summary of values from blood analytes and weight. This table provides the mean, standard deviation and p-values from the t-tests for each of the blood chemistry analytes and weight. The plus sign (+) indicates the values for individuals with plastic. The minus sign (-) indicates the values for individuals without plastic. The blood analytes measured were sodium (Na mmol/L), potassium (K mmol/L), chloride (Cl mmol/L), ionized calcium (iCa mmol/L), total carbon dioxide (TCO2), glucose (Glu mg/dL), Urea nitrogen/urea (BUN mg/dL), creatinine (Crea mg/dL), hematocrit (Hct %PCU), hemoglobin (Hb g/dL), anion gap (AnGap mmol/L) and are ordered respectively in the table. Differences were considered statistically significant when p < 0.07.

**Fig. S3.**
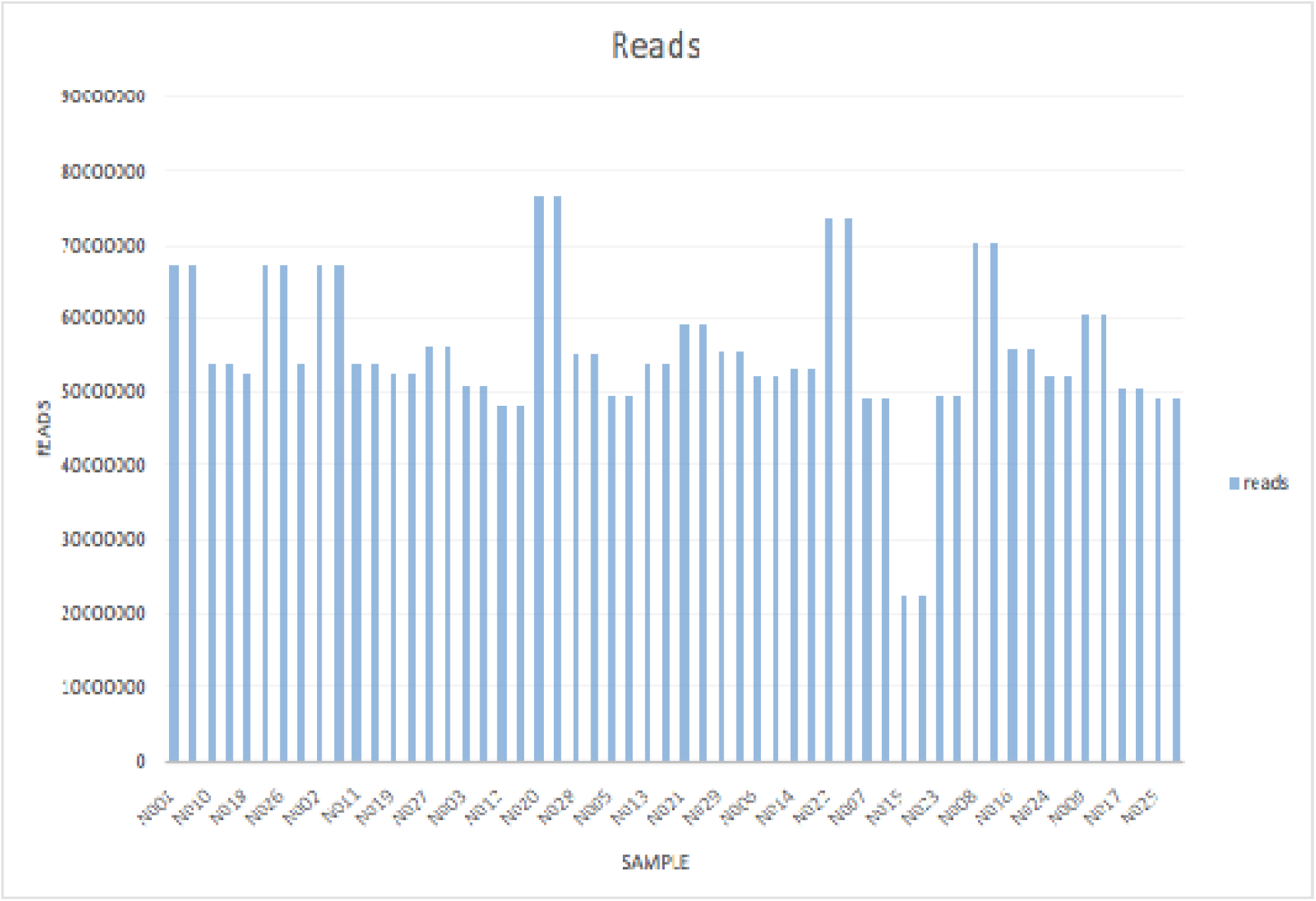
Reads per sample. This measurement assesses inferred sequence of base pairs that correspond to a single DNA fragment.

**Fig. S4.**
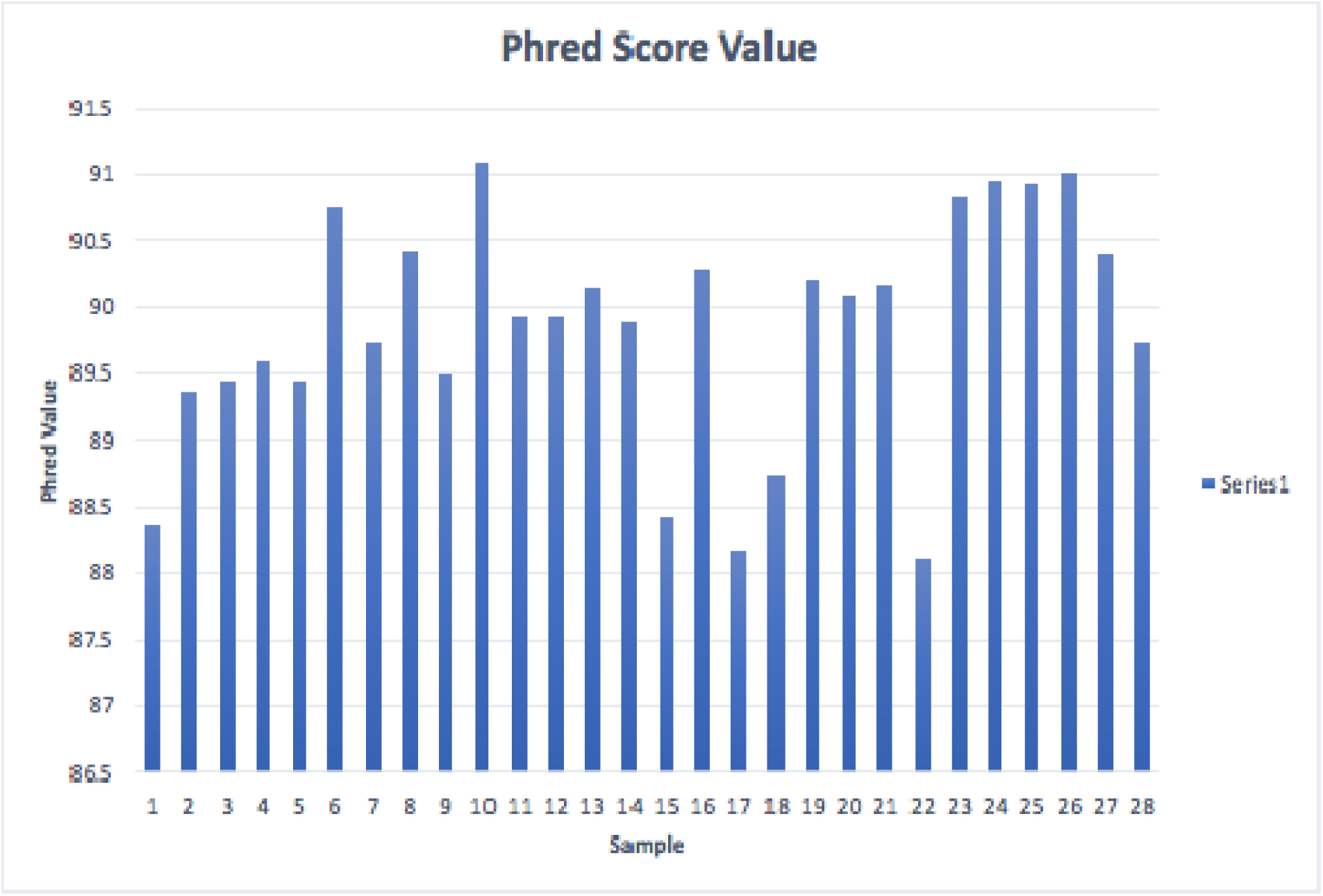
Phred score of samples. Phred score is used for quality assessment of sequencing

**Fig. S5.**
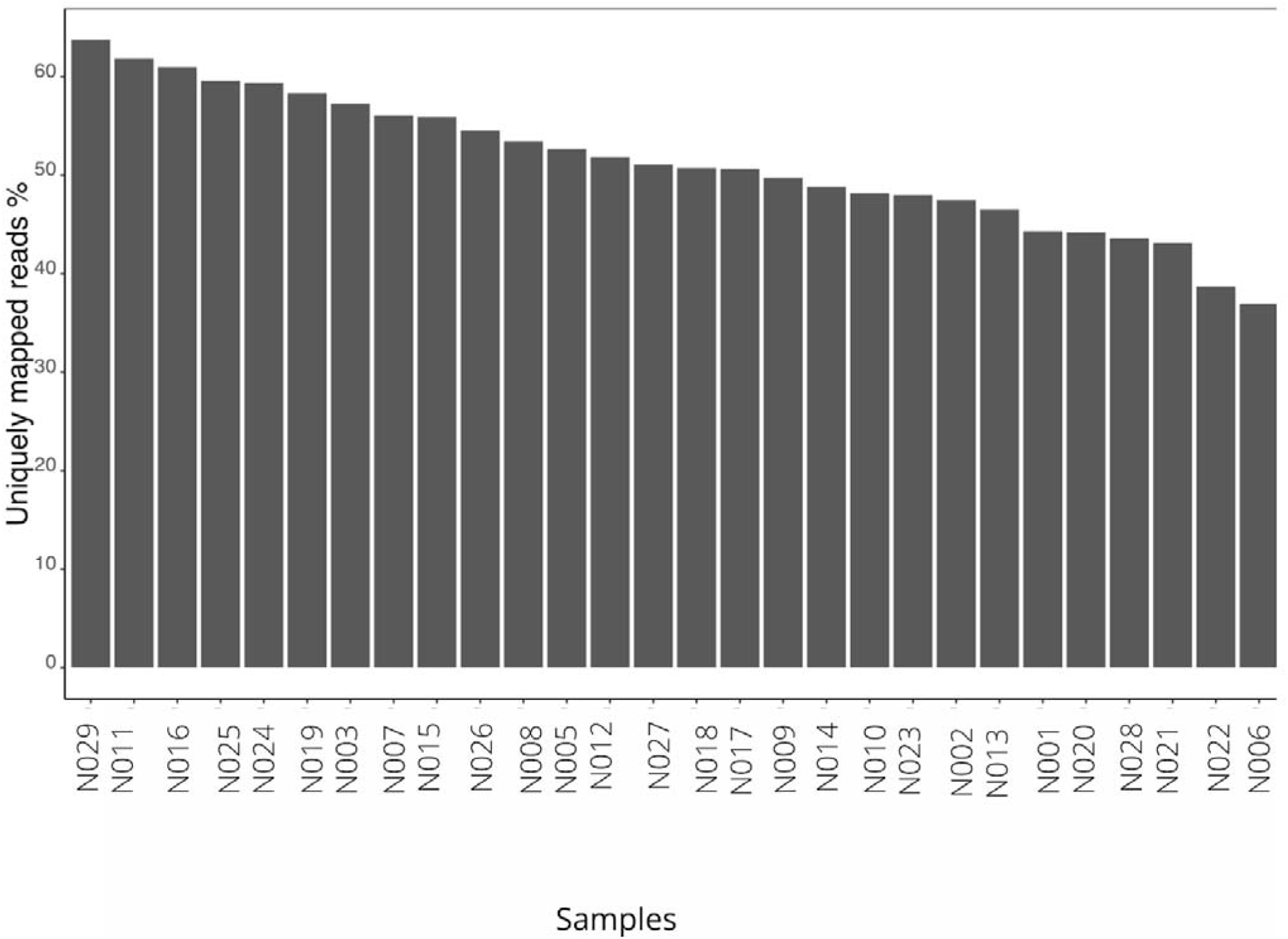
Uniquely mapped reads. Uniquely mapped reads have one exact location within the reference genome which they map to. This is the number of uniquely mapped reads from the prepared library that are aligned to the Cory Shearwater reference genome.

**Fig. S6.**
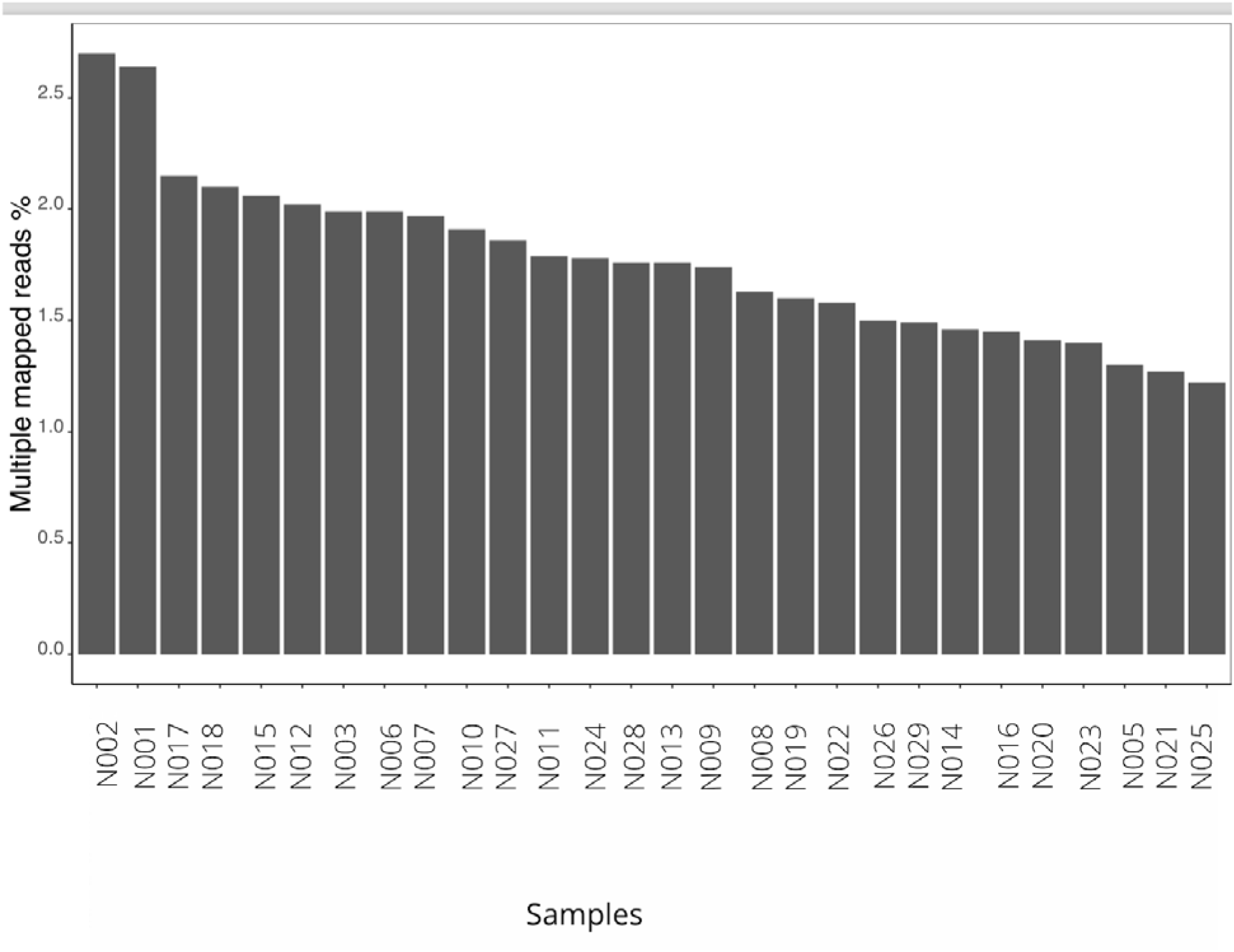
Multiply mapped reads. Multiple mapped reads are reads that map more than once in the genome. This is the number of multiple mapped reads from the prepared library that are aligned to the Cory Shearwater reference genome.

**Fig. S7.**
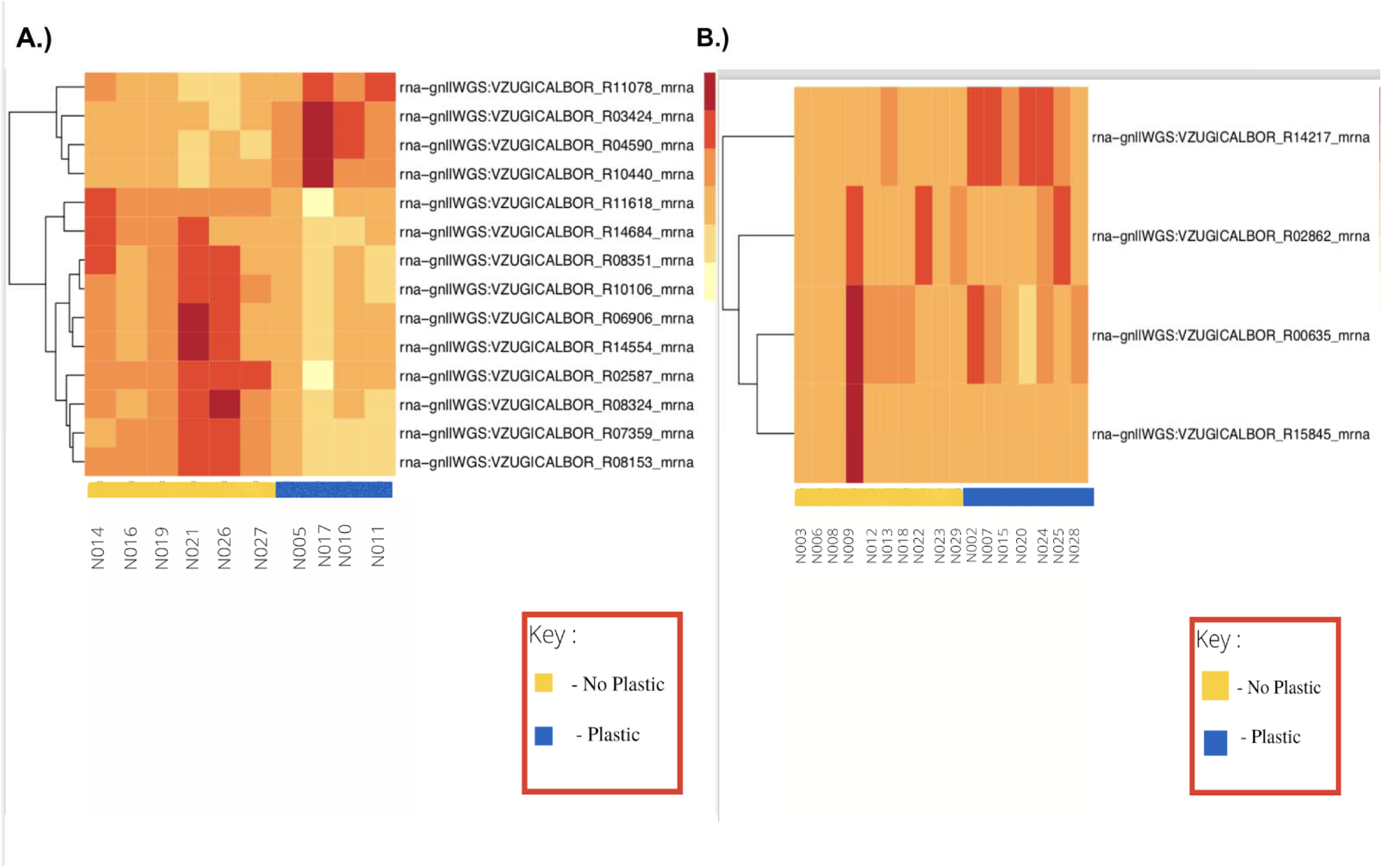
Differential gene expression analysis for plastic and sex. (A) Heatmap showing the 14 significantly differentially expressed genes in males. (B) Heatmap showing the 4 significant differentially expressed genes in females.

**Fig. S8.**
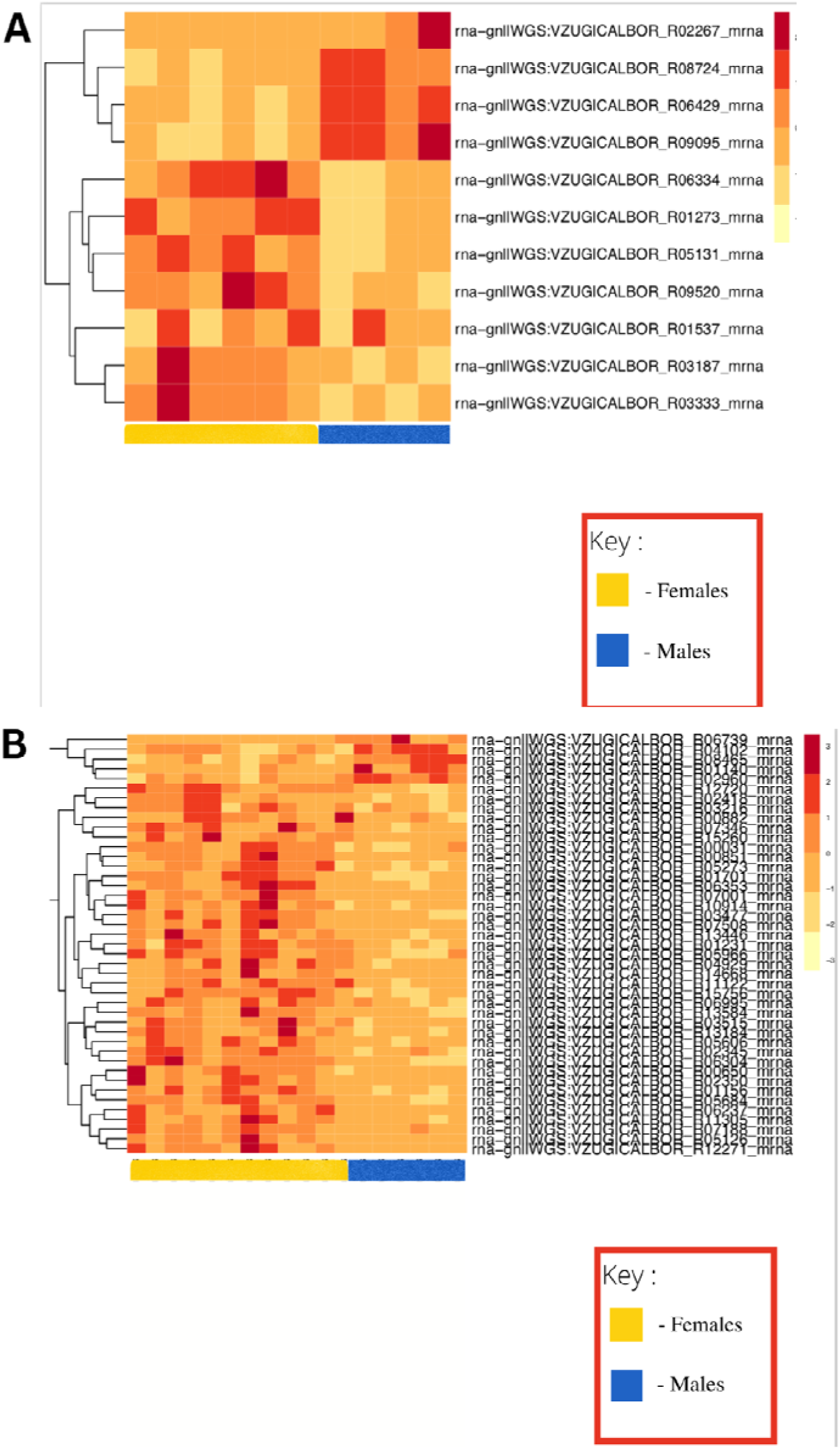
Differential gene expression analysis divided into samples that had ingested plastic and samples that did not have plastic. (A) Heatmap showing the 11 significant differentially expressed genes in all of the samples containing plastic separated by sex. (B) Heatmap showing the 43 significant differentially expressed genes in all samples that do not contain plastic separated by sex.

**Fig. S9.**
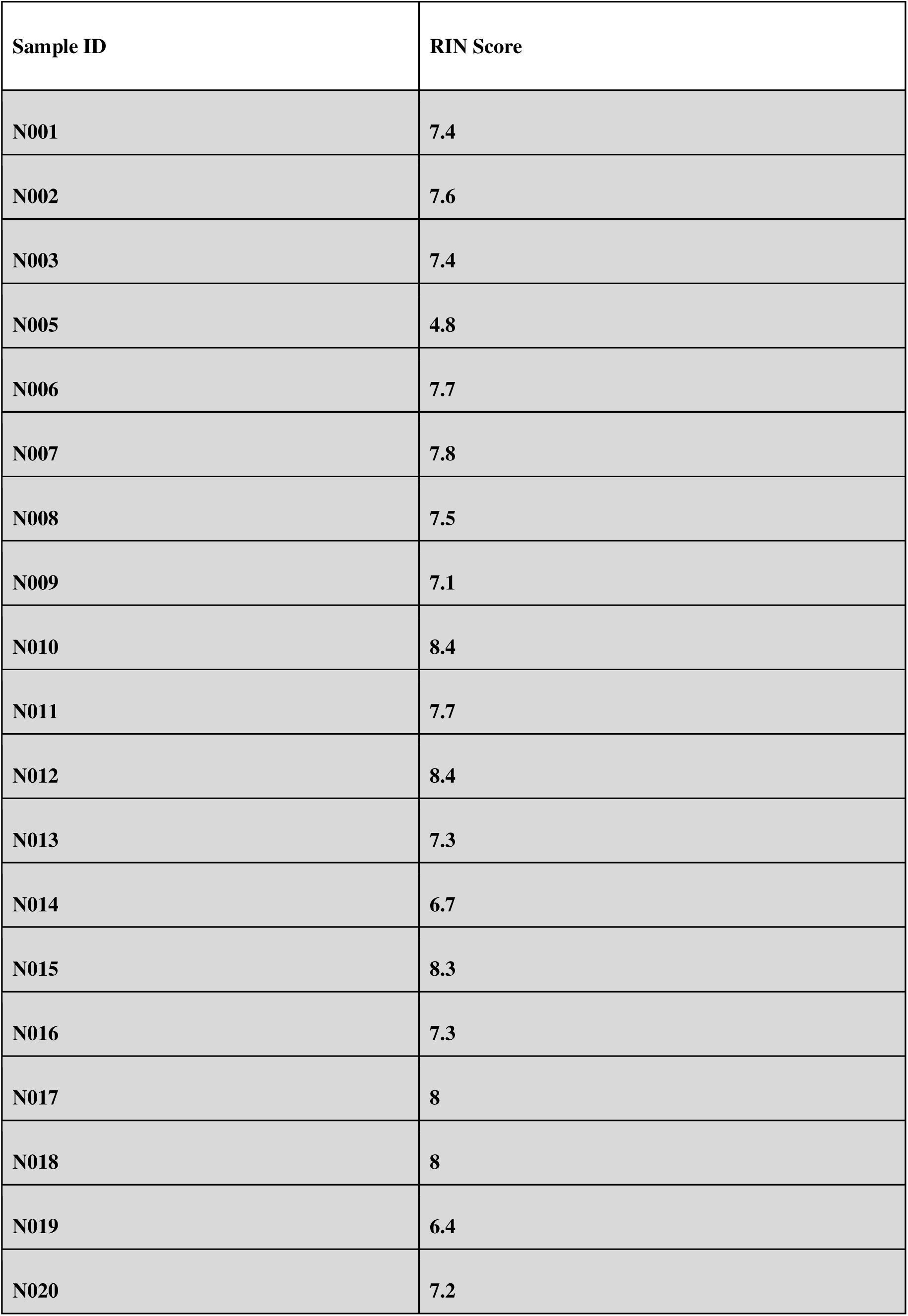
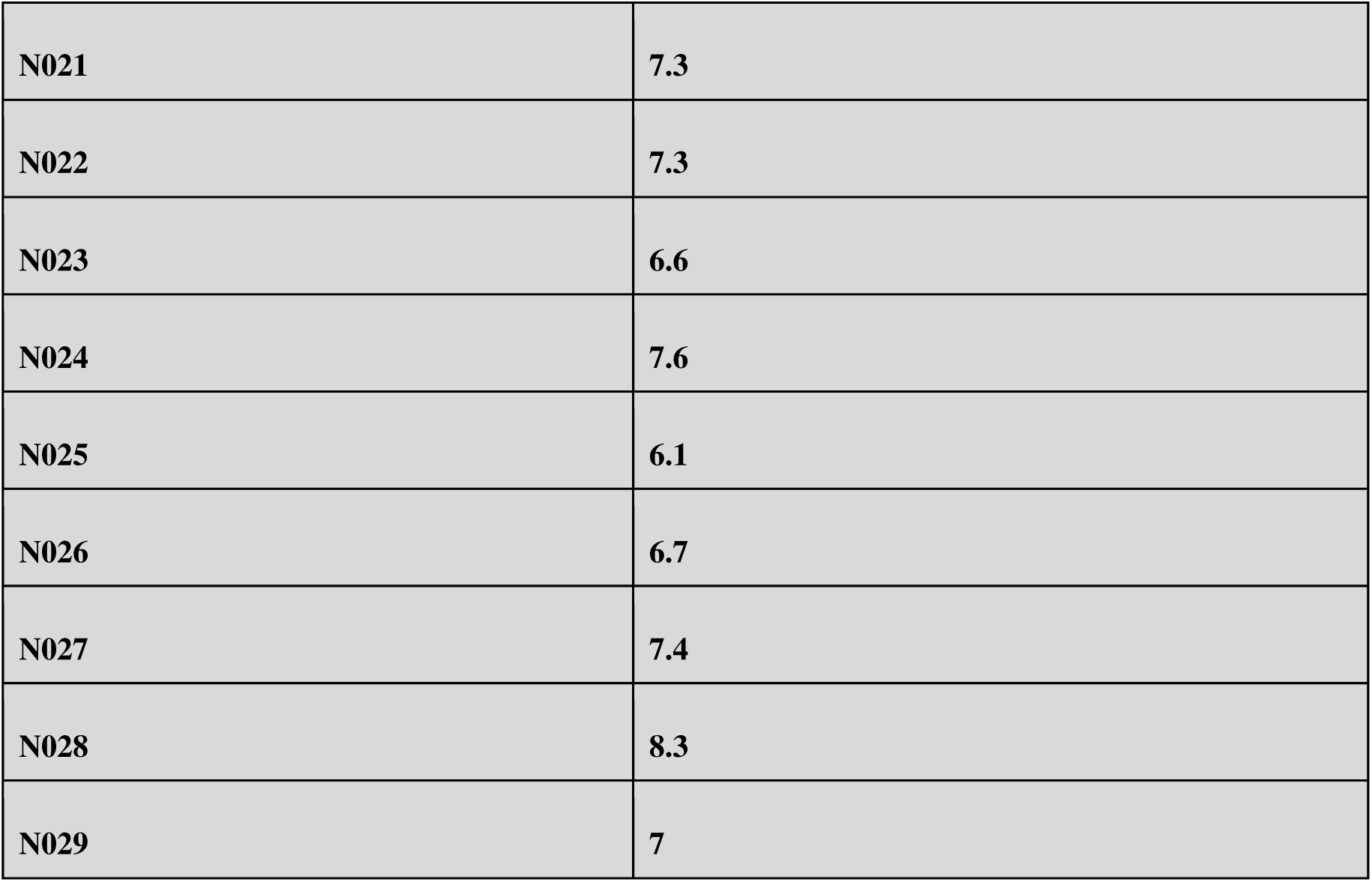
RIN Values. RIN values assign a numerical value to the quality of the RNA that we worked with. A RIN value of 8 and above indicated higher quality and integrity of RNA and values below 5 indicated some levels of RNA degradations.

**Fig. S10.**
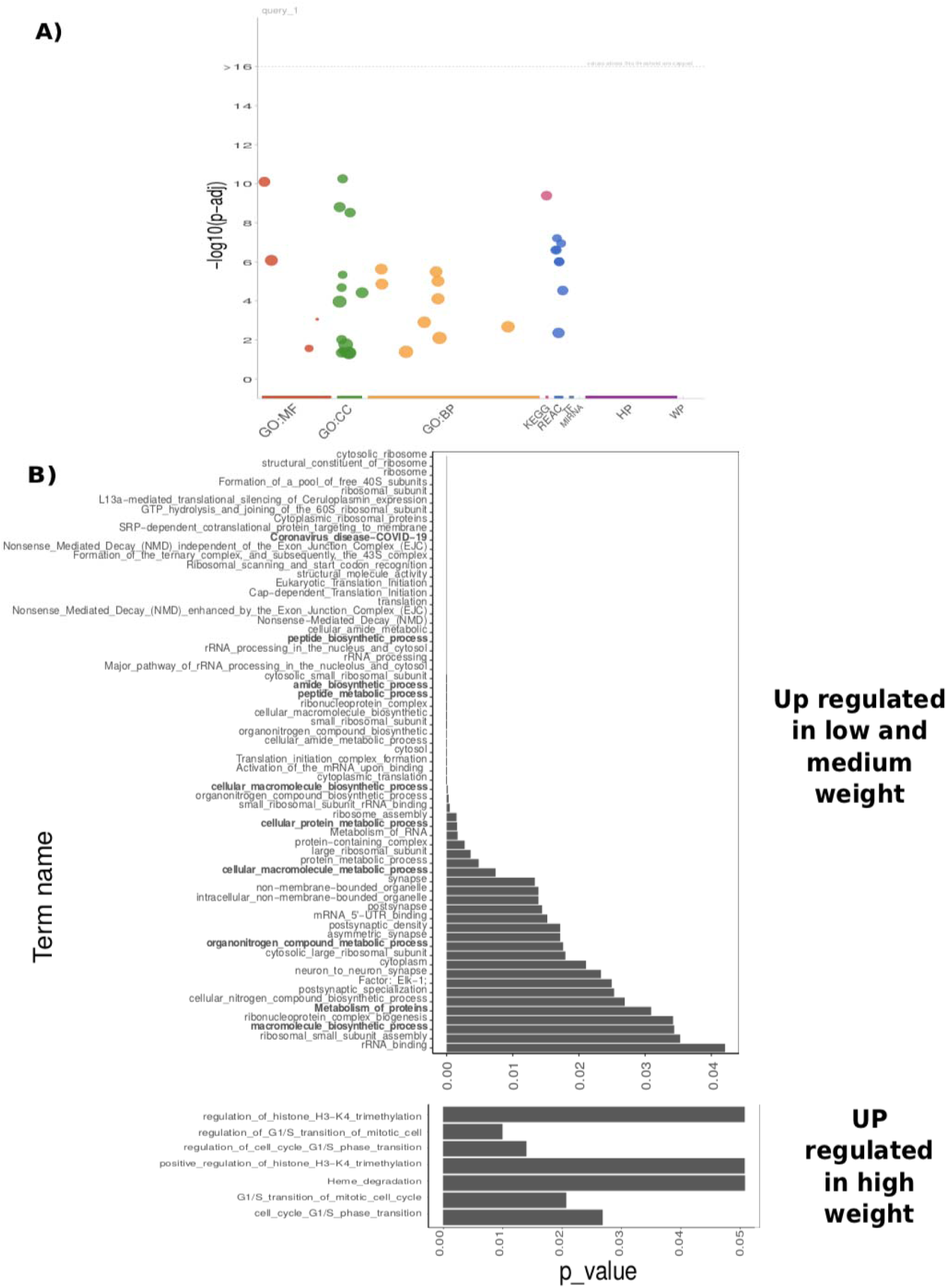
Gene enrichment analysis results for bird weight. (A) Manhattan plot showing results of enrichment analysis and the databases used. (B) Terms with significant values from the enrichment analysis in the differentially expressed genes between weight categories.

## References

Adams J, Felis J, Czapanskiy M. 2020. “Habitat Affinities and At-Sea Ranging Behaviors among Main Hawaiian Island Seabirds: Breeding Seabird Telemetry, 2013–2016.” BOEM

Barnes DKA, Galgani F, Thompson RC, Barlaz M. 2009. Accumulation and fragmentation of plastic debris in global environments. Philosophical Transactions of the Royal Society of London. Series B, Biological Sciences 364:1985–1998. DOI: 10.1098/rstb.2008.0205.

Barrett R, Camphuysen K. 2008. Sampling methods for seabird dietary studies. ICES Journal of Marine Science.

Barry ME, Pinto-González D, Orson FM, McKenzie GJ, Petry GR, Barry MA. 1999. Role of endogenous endonucleases and tissue site in transfection and CpG-mediated immune activation after naked DNA injection. Human Gene Therapy 10:2461–2480. DOI: 10.1089/10430349950016816.

Basu M, Wang K, Ruppin E, Hannenhalli S. 2021. Predicting tissue-specific gene expression from whole blood transcriptome. Science Advances 7:eabd6991. DOI: 10.1126/sciadv.abd6991.

Bittner GD, Yang CZ, Stoner MA. 2014. Estrogenic chemicals often leach from BPA-free plastic products that are replacements for BPA-containing polycarbonate products. Environmental Health 13:41. DOI: 10.1186/1476-069X-13-41.

Blumenberg M, Blumenberg M. 2019. Introductory Chapter. IntechOpen. DOI: 10.5772/intechopen.85980.

Boersma D, Groom M. 1993. Conservation of storm-petrels in the North Pacific. University of Washington.

Bonanno G, Orlando-Bonaca M. 2018. Ten inconvenient questions about plastics in the sea. Environmental Science & Policy 85:146–154. DOI: 10.1016/j.envsci.2018.04.005.

Bond AL, Lavers JL. 2013. Effectiveness of emetics to study plastic ingestion by Leach’s Storm-petrels (Oceanodroma leucorhoa). Marine Pollution Bulletin 70:171–175. DOI: 10.1016/j.marpolbul.2013.02.030.

Borrelle SB, Ringma J, Law KL, Monnahan CC, Lebreton L, McGivern A, Murphy E, Jambeck J, Leonard GH, Hilleary MA, Eriksen M, Possingham HP, De Frond H, Gerber LR, Polidoro B, Tahir A, Bernard M, Mallos N, Barnes M, Rochman CM. 2020. Predicted growth in plastic waste exceeds efforts to mitigate plastic pollution. Science 369:1515–1518. DOI: 10.1126/science.aba3656.

Bost, C. A., and LeMaho, Y.: 1993, ‘Seabirds as bio-indicators of changing marine ecosystems: new perspectives’, Acta Oecologica 14, 463–470.

Browne MA, Niven SJ, Galloway TS, Rowland SJ, Thompson RC. 2013. Microplastic Moves Pollutants and Additives to Worms, Reducing Functions Linked to Health and Biodiversity. Current Biology 23:2388–2392. DOI: 10.1016/j.cub.2013.10.012.

da Costa Araújo AP, de Melo NFS, de Oliveira Junior AG, Rodrigues FP, Fernandes T, de Andrade Vieira JE, Rocha TL, Malafaia G. 2020. How much are microplastics harmful to the health of amphibians? A study with pristine polyethylene microplastics and Physalaemus cuvieri. Journal of Hazardous Materials 382:121066. DOI: 10.1016/j.jhazmat.2019.121066.

Cózar A, Echevarría F, González-Gordillo JI, Irigoien X, Úbeda B, Hernández-León S, Palma ÁT, Navarro S, García-de-Lomas J, Ruiz A, Fernández-de-Puelles ML, Duarte CM. 2014. Plastic debris in the open ocean. Proceedings of the National Academy of Sciences 111:10239–10244. DOI: 10.1073/pnas.1314705111.

DNA Microarray Technology Fact Sheet. Available at https://www.genome.gov/about-genomics/fact-sheets/DNA-Microarray-Technology (accessed November 8, 2022).

Dobin A, Davis CA, Schlesinger F, Drenkow J, Zaleski C, Jha S, Batut P, Chaisson M, Gingeras TR. 2013. STAR: ultrafast universal RNA-seq aligner. Bioinformatics (Oxford, England) 29:15–21. DOI: 10.1093/bioinformatics/bts635.

Dobin A, Gingeras TR. 2015. Mapping RNA-seq Reads with STAR. Current Protocols in Bioinformatics 51:11.14.1–11.14.19. DOI: 10.1002/0471250953.bi1114s51.

Duffy DC, Jackson S. 1986. Diet Studies of Seabirds: A Review of Methods. Colonial Waterbirds 9:1–17. DOI: 10.2307/1521138.

Ekblom R, Wennekes P, Horsburgh GJ, Burke T. 2014. Characterization of the house sparrow (Passer domesticus) transcriptome: a resource for molecular ecology and immunogenetics. Molecular Ecology Resources 14:636–646. DOI: 10.1111/1755-0998.12213.

Estrogen’s Effects on the Female Body. 2022. *Available at* https://www.hopkinsmedicine.org/health/conditions-and-diseases/estrogens-effects-on-the-female-body (accessed November 8, 2022).

Feng S. et al. Dense sampling of bird diversity increases power of comparative genomics. Nature 587:252–257. DOI: 10.1038/s41586-020-2873-9.

Furness R. W., Monaghan P. 1987. Seabird Ecology. Blackie; Published in the USA by Chapman and Hall.

Franchini P, Irisarri I, Fudickar A, Schmidt A, Meyer A, Wikelski M, Partecke J. 2017. Animal tracking meets migration genomics: transcriptomic analysis of a partially migratory bird species. Molecular Ecology 26:3204–3216. DOI: 10.1111/mec.14108.

Fridolfsson A-K, Ellegren H. 1999. A Simple and Universal Method for Molecular Sexing of Non-Ratite Birds. Journal of Avian Biology 30:116–121. DOI: 10.2307/3677252.

Laist DW. 1997. Impacts of Marine Debris: Entanglement of Marine Life in Marine Debris Including a Comprehensive List of Species with Entanglement and Ingestion Records. In: Coe JM, Rogers DB eds. *Marine Debris: Sources, Impacts, and Solutions*. Springer Series on Environmental Management. New York, NY: Springer, 99–139. DOI: 10.1007/978-1-4613-8486-1_10.

The Problem of Marine Plastic Pollution. 2016. *Available at* https://www.cleanwater.org/problem-marine-plastic-pollution (accessed November 8, 2022).

Clinical Pathology of Plastic Ingestion in Marine Birds and Relationships with Blood Chemistry | Environmental Science & Technology. Available at https://pubs.acs.org/doi/full/10.1021/acs.est.9b02098 (accessed November 8, 2022).

Coe JM, Rogers DB. 1997. Introduction. In: Coe JM, Rogers DB eds. Marine Debris: Sources, Impacts, and Solutions. Springer Series on Environmental Management. New York, NY: Springer, 5–6. DOI: 10.1007/978-1-4613-8486-1_1.

Fine-scale population genetic structure and barriers to gene flow in a widespread seabird (Ardenna pacifica) | Biological Journal of the Linnean Society | Oxford Academic. Available at https://academic.oup.com/biolinnean/article/137/1/125/6650017 (accessed November 8, 2022).

Gall SC, Thompson RC. 2015. The impact of debris on marine life. Marine Pollution Bulletin 92:170–179. DOI: 10.1016/j.marpolbul.2014.12.041.

Gallo F, Fossi C, Weber R, Santillo D, Sousa J, Ingram I, Nadal A, Romano D. 2018. Marine litter plastics and microplastics and their toxic chemicals components: the need for urgent preventive measures. Environmental Sciences Europe 30:13. DOI: 10.1186/s12302-018-0139-z.

Gao H, Yang B-J, Li N, Feng L-M, Shi X-Y, Zhao W-H, Liu S-J. 2015. Bisphenol A and Hormone-Associated Cancers: Current Progress and Perspectives. Medicine 94:e211. DOI: 10.1097/MD.0000000000000211.

Gornicka A, Morris-Stiff G, Thapaliya S, Papouchado BG, Berk M, Feldstein AE. 2011. Transcriptional Profile of Genes Involved in Oxidative Stress and Antioxidant Defense in a Dietary Murine Model of Steatohepatitis. Antioxidants & Redox Signaling 15:437–445. DOI: 10.1089/ars.2010.3815.

gprofiler2 -- an R package for gene list functional… | F1000Research. *Available at* https://f1000research.com/articles/9-709/v2 (accessed November 8, 2022).

Handling multi-mapped reads — seqcluster 1.2.4a7 documentation. *Available at* https://seqcluster.readthedocs.io/multi_mapped.html (accessed November 8, 2022).

Harr KE. 2002. Clinical Chemistry of Companion Avian Species: A Review. Veterinary Clinical Pathology 31:140–151. DOI: 10.1111/j.1939-165X.2002.tb00295.x.

Harrison AG, Lin T, Wang P. 2020. Mechanisms of SARS-CoV-2 Transmission and Pathogenesis. Trends in Immunology 41:1100–1115. DOI: 10.1016/j.it.2020.10.004.

Herman RW, Winger BM, Dittmann DL, Harvey MG. 2022. Fine-scale population genetic structure and barriers to gene flow in a widespread seabird (Ardenna pacifica). Biological Journal of the Linnean Society 137:125–136. DOI: 10.1093/biolinnean/blac091.

Hirt N, Body-Malapel M. 2020. Immunotoxicity and intestinal effects of nano- and microplastics: a review of the literature. Particle and Fibre Toxicology 17:57. DOI: 10.1186/s12989-020-00387-7.

Hochleithner. “Biochemistries.” Avian Medicine: Principles and Applications, pp. 243–245

Jâms IB, Windsor FM, Poudevigne-Durance T, Ormerod SJ, Durance I. 2020. Estimating the size distribution of plastics ingested by animals. Nature Communications 11:1594. DOI: 10.1038/s41467-020-15406-6.

Jax E, Müller I, Börno S, Borlinghaus H, Eriksson G, Fricke E, Timmermann B, Pendl H, Fiedler W, Klein K, Schreiber F, Wikelski M, Magor KE, Kraus RHS. 2021. Health monitoring in birds using bio-loggers and whole blood transcriptomics. Scientific Reports 11:10815. DOI: 10.1038/s41598-021-90212-8.

Jax E, Wink M, Kraus RHS. 2018. Avian transcriptomics: opportunities and challenges. Journal of Ornithology 159:599–629. DOI: 10.1007/s10336-018-1532-5.

Kain EC, Lavers JL, Berg CJ, Raine AF, Bond AL. 2016. Plastic ingestion by Newell’s (Puffinus newelli) and wedge-tailed shearwaters (Ardenna pacifica) in Hawaii. Environmental Science and Pollution Research 23:23951–23958. DOI: 10.1007/s11356-016-7613-1.

Karami A, Romano N, Galloway T, Hamzah H. 2016. Virgin microplastics cause toxicity and modulate the impacts of phenanthrene on biomarker responses in African catfish (Clarias gariepinus). Environmental Research 151:58–70. DOI: 10.1016/j.envres.2016.07.024.

Kenyon KW, Kridler E. 1969. Laysan Albatrosses swallow indigestible matter. The Auk 86:339– 343. DOI: 10.2307/4083505.

Koelmans AA, Bakir A, Burton GA, Janssen CR. 2016. Microplastic as a Vector for Chemicals in the Aquatic Environment: Critical Review and Model-Supported Reinterpretation of Empirical Studies. Environmental Science & Technology 50:3315–3326. DOI: 10.1021/acs.est.5b06069.

Krjutškov K, Koel M, Roost AM, Katayama S, Einarsdottir E, Jouhilahti E-M, Söderhäll C, Jaakma Ü, Plaas M, Vesterlund L, Lohi H, Salumets A, Kere J. 2016. Globin mRNA reduction for whole-blood transcriptome sequencing. Scientific Reports 6:31584. DOI: 10.1038/srep31584.

Labocha MK, Hayes JP. 2012. Morphometric indices of body condition in birds: a review. Journal of Ornithology 153:1–22. DOI: 10.1007/s10336-011-0706-1.

Laing LV, Viana J, Dempster EL, Trznadel M, Trunkfield LA, Uren Webster TM, van Aerle R, Paull GC, Wilson RJ, Mill J, Santos EM. 2016. Bisphenol A causes reproductive toxicity, decreases dnmt1 transcription, and reduces global DNA methylation in breeding zebrafish (Danio rerio). Epigenetics 11:526–538. DOI: 10.1080/15592294.2016.1182272.

Li B., Dewey C.N. 2021.“RSEM: accurate transcript quantification from RNA-Seq data with or without a reference genome”. BMC Bioinformatics 12, 323.

Limonta G, Mancia A, Benkhalqui A, Bertolucci C, Abelli L, Fossi MC, Panti C. 2019. Microplastics induce transcriptional changes, immune response and behavioral alterations in adult zebrafish. Scientific Reports 9:15775. DOI: 10.1038/s41598-019-52292-5.

Love MI, Huber W, Anders S. 2014. Moderated estimation of fold change and dispersion for RNA-seq data with DESeq2. Genome Biology 15:550. DOI: 10.1186/s13059-014-0550-8.

Lu Y, Zhang Y, Deng Y, Jiang W, Zhao Y, Geng J, Ding L, Ren H. 2016. Uptake and Accumulation of Polystyrene Microplastics in Zebrafish (Danio rerio) and Toxic Effects in Liver. Environmental Science & Technology 50:4054–4060. DOI: 10.1021/acs.est.6b00183.

Mallory ML, Robinson SA, Hebert CE, Forbes MR. 2010. Seabirds as indicators of aquatic ecosystem conditions: A case for gathering multiple proxies of seabird health. Marine Pollution Bulletin 60:7–12. DOI: 10.1016/j.marpolbul.2009.08.024.

Meitern R, Andreson R, Hõrak P. 2014. Profile of whole blood gene expression following immune stimulation in a wild passerine. BMC Genomics 15:533. DOI: 10.1186/1471-2164-15-533.

Microplastics | National Geographic Society. Available at https://education.nationalgeographic.org/resource/microplastics (accessed November 8, 2022).

Mootha VK, Lindgren CM, Eriksson K-F, Subramanian A, Sihag S, Lehar J, Puigserver P, Carlsson E, Ridderstråle M, Laurila E, Houstis N, Daly MJ, Patterson N, Mesirov JP, Golub TR, Tamayo P, Spiegelman B, Lander ES, Hirschhorn JN, Altshuler D, Groop LC. 2003. PGC-1α -responsive genes involved in oxidative phosphorylation are coordinately downregulated in human diabetes. Nature Genetics 34:267–273. DOI: 10.1038/ng1180.

Morphometric indices of body condition in birds: a review | SpringerLink. Available at https://link.springer.com/article/10.1007/s10336-011-0706-1 (accessed November 8, 2022).

Nania TG, Shugart GW. 2021. Are plastic particles reduced in size in seabirds’ stomachs? Marine Pollution Bulletin 172:112843. DOI: 10.1016/j.marpolbul.2021.112843.

Peng Y, Wu P, Schartup AT, Zhang Y. 2021. Plastic waste release caused by COVID-19 and its fate in the global ocean. Proceedings of the National Academy of Sciences 118:e2111530118. DOI: 10.1073/pnas.2111530118.

Pierce KE, Harris RJ, Larned L, Pokras M. 2004. Obstruction and starvation associated with plastic ingestion in a Northern Gannet Morus bassanus and a Greater Shearwater Puffinus gravis. Marine Ornithology 32:187–189.

Prata JC, da Costa JP, Lopes I, Duarte AC, Rocha-Santos T. 2020. Environmental exposure to microplastics: An overview on possible human health effects. Science of The Total Environment 702:134455. DOI: 10.1016/j.scitotenv.2019.134455.

Provencher JF, Borrelle SB, Bond AL, Lavers JL, van Franeker JA, Kühn S, Hammer S, Avery-Gomm S, Mallory ML. 2019. Recommended best practices for plastic and litter ingestion studies in marine birds: Collection, processing, and reporting. FACETS 4:111–130. DOI: 10.1139/facets-2018-0043.

Puchta M, Boczkowska M, Groszyk J. 2020. Low RIN Value for RNA-Seq Library Construction from Long-Term Stored Seeds: A Case Study of Barley Seeds. Genes 11:1190. DOI: 10.3390/genes11101190.

Ritchie H, Roser M. 2018. Plastic Pollution. Our World in Data.

Ryan PG. 1987. The effects of ingested plastic on seabirds: Correlations between plastic load and body condition. Environmental Pollution 46:119–125. DOI: 10.1016/0269-7491(87)90197-7.

Scholz M. 2021. “Metagenomics - Phred score (Q score).” Metagenomics wiki, https://www.metagenomics.wiki/pdf/definition/phred-base-quality.

Science-Based Solutions to Plastic Pollution. 2020. One Earth 2:5–7. DOI: 10.1016/j.oneear.2020.01.00.

Sievert P.R., Sileo L. 1993. “The effects of ingested plastic on growth and survival of albatross chicks. In: Vermeer, K., Briggs, K.T., Morgan, K.H. & Siegal–Causey, D. (Eds). The status, ecology, and conservation of marine birds of the North Pacific.” Ottawa: Canadian Wildlife Service Special Publication, pp. 212–217.

Signa G, Mazzola A, Vizzini S. 2021. Seabird influence on ecological processes in coastal marine ecosystems: An overlooked role? A critical review. Estuarine, Coastal and Shelf Science 250:107164. DOI: 10.1016/j.ecss.2020.107164.

Stafford R, Jones PJS. 2019. Viewpoint – Ocean plastic pollution: A convenient but distracting truth? Marine Policy 103:187–191. DOI: 10.1016/j.marpol.2019.02.003.

Sun J, Fang R, Wang H, Xu D-X, Yang J, Huang X, Cozzolino D, Fang M, Huang Y. 2022. A review of environmental metabolism disrupting chemicals and effect biomarkers associating disease risks: Where exposomics meets metabolomics. Environment International 158:106941. DOI: 10.1016/j.envint.2021.106941.

Tanaka K, Takada H, Yamashita R, Mizukawa K, Fukuwaka M, Watanuki Y. 2013. Accumulation of plastic-derived chemicals in tissues of seabirds ingesting marine plastics. Marine Pollution Bulletin 69:219–222. DOI: 10.1016/j.marpolbul.2012.12.010.

“Terms and Definitions – ENCODE.” *encode*, https://www.encodeproject.org/data-standards/terms/.

Tilman D. 1989. Ecological Experimentation: Strengths and Conceptual Problems. In: Likens GE ed. Long-Term Studies in Ecology: Approaches and Alternatives. New York, NY: Springer, 136–157. DOI: 10.1007/978-1-4615-7358-6_6.

Valle CA, Ulloa C, Regalado C, Muñoz-Pérez J-P, Garcia J, Hardesty BD, Skehel A, Deresienski D, Lewbart GA. 2020. HEALTH STATUS AND BASELINE HEMATOLOGY, BIOCHEMISTRY, AND BLOOD GAS VALUES OF GALAPAGOS SHEARWATERS (PUFFINUS SUBALARIS). Journal of Zoo and Wildlife Medicine 50:1026–1030. DOI: 10.1638/2019-0035R.

Verla AW, Enyoh CE, Verla EN, Nwarnorh KO. 2019. Microplastic–toxic chemical interaction: a review study on quantified levels, mechanism and implication. SN Applied Sciences 1:1400. DOI: 10.1007/s42452-019-1352-0.

Verlis KM, Campbell ML, Wilson SP. 2018. Seabirds and plastics don’t mix: Examining the differences in marine plastic ingestion in wedge-tailed shearwater chicks at near-shore and offshore locations. Marine Pollution Bulletin 135:852–861. DOI: 10.1016/j.marpolbul.2018.08.016.

Wang L, Nabi G, Yin L, Wang Y, Li S, Hao Z, Li D. 2021. Birds and plastic pollution: recent advances. Avian Research 12:59. DOI: 10.1186/s40657-021-00293-2.

Weimerskirch H, Grissac S de, Ravache A, Prudor A, Corbeau A, Congdon BC, McDuie F, Bourgeois K, Dromzée S, Butscher J, Menkes C, Allain V, Vidal E, Jaeger A, Borsa P. 2020. At-sea movements of wedge-tailed shearwaters during and outside the breeding season from four colonies in New Caledonia. Marine Ecology Progress Series 633:225–238. DOI: 10.3354/meps13171.

Wilcox C, Van Sebille E, Hardesty BD. 2015. Threat of plastic pollution to seabirds is global, pervasive, and increasing. Proceedings of the National Academy of Sciences 112:11899–11904. DOI: 10.1073/pnas.1502108112.

Xu K, Zhang Y, Huang Y, Wang J. 2021. Toxicological effects of microplastics and phenanthrene to zebrafish (Danio rerio). Science of The Total Environment 757:143730. DOI: 10.1016/j.scitotenv.2020.143730.

## Supporting References

Ashraf MA. 2017. Persistent organic pollutants (POPs): a global issue, a global challenge. Environmental Science and Pollution Research 24:4223–4227. DOI: 10.1007/s11356-015-5225-9.

Bighiu MA, Gorokhova E, Almroth BC, Wiklund A-KE. 2017. Metal contamination in harbours impacts life-history traits and metallothionein levels in snails. PLOS ONE 12:e0180157. DOI:10.1371/journal.pone.0180157.

Briffa J, Sinagra E, Blundell R. 2020. Heavy metal pollution in the environment and their toxicological effects on humans. Heliyon 6:e04691. DOI: 10.1016/j.heliyon.2020.e04691.

Claessens M, Meester SD, Landuyt LV, Clerck KD, Janssen CR. 2011. Occurrence and distribution of microplastics in marine sediments along the Belgian coast. Marine Pollution Bulletin 62:2199–2204. DOI: 10.1016/j.marpolbul.2011.06.030.

Désert C, Merlot E, Zerjal T, Bed’hom B, Härtle S, Le Cam A, Roux P-F, Baeza E, Gondret F, Duclos MJ, Lagarrigue S. 2016. Transcriptomes of whole blood and PBMC in chickens. Comparative Biochemistry and Physiology Part D: Genomics and Proteomics 20:1–9. DOI:10.1016/j.cbd.2016.06.008.

Liu S, Shi J, Wang J, Dai Y, Li H, Li J, Liu X, Chen X, Wang Z, Zhang P. 2021. Interactions Between Microplastics and Heavy Metals in Aquatic Environments: A Review. Frontiers in Microbiology 12:652520. DOI: 10.3389/fmicb.2021.652520.

Liu J, Walter E, Stenger D, Thach D. 2006. Effects of Globin mRNA Reduction Methods on Gene Expression Profiles from Whole Blood. The Journal of Molecular Diagnostics 8:551–558. DOI: 10.2353/jmoldx.2006.060021.

Lobenhofer EK, Auman JT, Blackshear PE, Boorman GA, Bushel PR, Cunningham ML, Fostel JM, Gerrish K, Heinloth AN, Irwin RD, Malarkey DE, Merrick BA, Sieber SO, Tucker CJ, Ward SM, Wilson RE, Hurban P, Tennant RW, Paules RS. 2008. Gene expression response in target organ and whole blood varies as a function of target organ injury phenotype. Genome Biology 9:R100. DOI: 10.1186/gb-2008-9-6-r100.

NOAA. “A Guide to Plastic in the Ocean.” NOAA’s National Ocean Service, https://oceanservice.noaa.gov/hazards/marinedebris/plastics-in-the-ocean.html.

NOAA. “Ocean Garbage Patches.” NOAA’s National Ocean Service, https://oceanservice.noaa.gov/podcast/mar18/nop14-ocean-garbage-patches.html.

Plastic debris in the open ocean | PNAS. *Available at* https://www.pnas.org/doi/full/10.1073/pnas.1314705111 (accessed November 8, 2022).

Shaffer SA, Tremblay Y, Weimerskirch H, Scott D, Thompson DR, Sagar PM, Moller H, Taylor GA, Foley DG, Block BA, Costa DP. 2006. Migratory shearwaters integrate oceanic resources across the Pacific Ocean in an endless summer. Proceedings of the National Academy of Sciences 103:12799–12802. DOI: 10.1073/pnas.0603715103.

